# Write and Read: Harnessing Synthetic DNA Modifications for Nanopore Sequencing

**DOI:** 10.1101/2025.04.14.648727

**Authors:** Uri Bertocchi, Assaf Grunwald, Gal Goldner, Eliran Eitan, Sigal Avraham, Shani Dvir, Jasline Deek, Yael Michaeli, Brian Yao, Jennifer Listgarten, Jared T. Simpson, Winston Timp, Yuval Ebenstein

## Abstract

An exciting feature of nanopore sequencing is its ability to record multi-omic information on the same sequenced DNA molecule. Well-trained models allow the detection of nucleotide-specific molecular signatures through changes in ionic current as DNA molecules translocate through the nanopore. Thus, naturally occurring DNA modifications, such as DNA methylation and hydroxymethylation, may be recorded simultaneously with the genetic sequence. Additional genomic information, such as chromatin state or the locations of bound transcription factors, may also be recorded if their locations are chemically encoded into the DNA. Here, we present a versatile “write-and-read” framework, where chemo-enzymatic DNA labeling with unnatural synthetic tags results in predictable electrical fingerprints in nanopore sequencing. As a proof-of-concept, we explore a DNA glucosylation approach that selectively modifies 5-hydroxymethylcytosine (5hmC) with glucose or glucose-azide adducts. We demonstrate that these modifications generate distinct and reproducible electrical shifts, enabling the direct detection of chemically altered nucleotides. We further demonstrate that enzymatic alkylation, such as the enzymatic transfer of azide residues to the N6 position of adenines, also produces characteristic nanopore signal shifts relative to the native adenine and 6-methyladenine. Beyond direct nucleotide detection, this approach introduces new possibilities for bio-orthogonal DNA labeling, enabling an extended alphabet of sequence-specific detectable moieties. The future use of programmable chemical modifications for simultaneous analysis of multiple omics features on individual molecules opens new avenues for genetic research and discovery.

---

Nanopore sequencing has emerged as a powerful platform in genomics, offering the unique capability to read long nucleic acid molecules at the single-molecule level^1–6^. Protein nanopores function as ultrasensitive molecular sensors, that detect changes in ionic current as individual molecules traverse the pore^6,7^. When an applied voltage drives ionic current through the nanopore, the passage of a molecular species attenuates the flow, producing a measurable change in current^6,7^. The modulation is influenced by the size of the molecule and its interaction with the pore’s interior through electrostatic and hydrophobic forces. These fundamental principles underlie nanopore-based sequencing, where the continuous monitoring of ionic current enables real-time profiling of single-stranded DNA^6^, RNA^8^, or proteins^9^, as they are translocated through the nanopore.

One of the advantages of protein nanopores, such as those offered by Oxford Nanopore Technologies (ONT) for sequencing, is that the electrical signals detected during the process are attenuated by chemical modifications of nucleobases^2^. This sensitivity makes nanopore sequencing especially valuable for epigenetic studies^1,2,10,11^. To date, commercial nanopore base callers have been trained primarily to detect naturally occurring modifications (e.g., 5- methylcytosine, N -methyladenine)^12^. However, the exact electrical readout can further be expanded to report on a far broader palette of synthetic labels introduced by chemistry.

In parallel to nanopore-based approaches, several in vitro reactions for DNA repair, glucosylation, and alkylation have been developed to attach tags to epigenetic and genetic markers^13–19^; these methods have been widely used for optical genome mapping to analyze long-range genetic and epigenetic profiles^20–30^. We hypothesized that the same approach may be applied to electrical detection by introducing chemical moieties that create electrical contrast and are distinctly detectable via nanopore sequencing.

This proof-of-concept study demonstrates selective chemoenzymatic labeling with various synthetic modifications for nanopore sequencing. These modifications affect the measured current in the nanopore, enabling specific detection of these tags. This approach enables precise control over which DNA targets are chemically marked, integrating three key components: (1) DNA-modifying enzymes that recognize specific sequence motifs or pre-existing modifications^31,32^, (2) engineered cofactors carrying detectable chemical groups^33–39^, and (3) high-resolution electrical detection of labeled DNA strands via nanopore sequencing; for an illustration, see Figure 1. Together, these elements establish a versatile “write-and-read” paradigm, wherein a chosen enzyme installs a tailored chemical marker at specific genomic sites, generating distinct current signatures in the nanopore readout.

**Figure 1.**
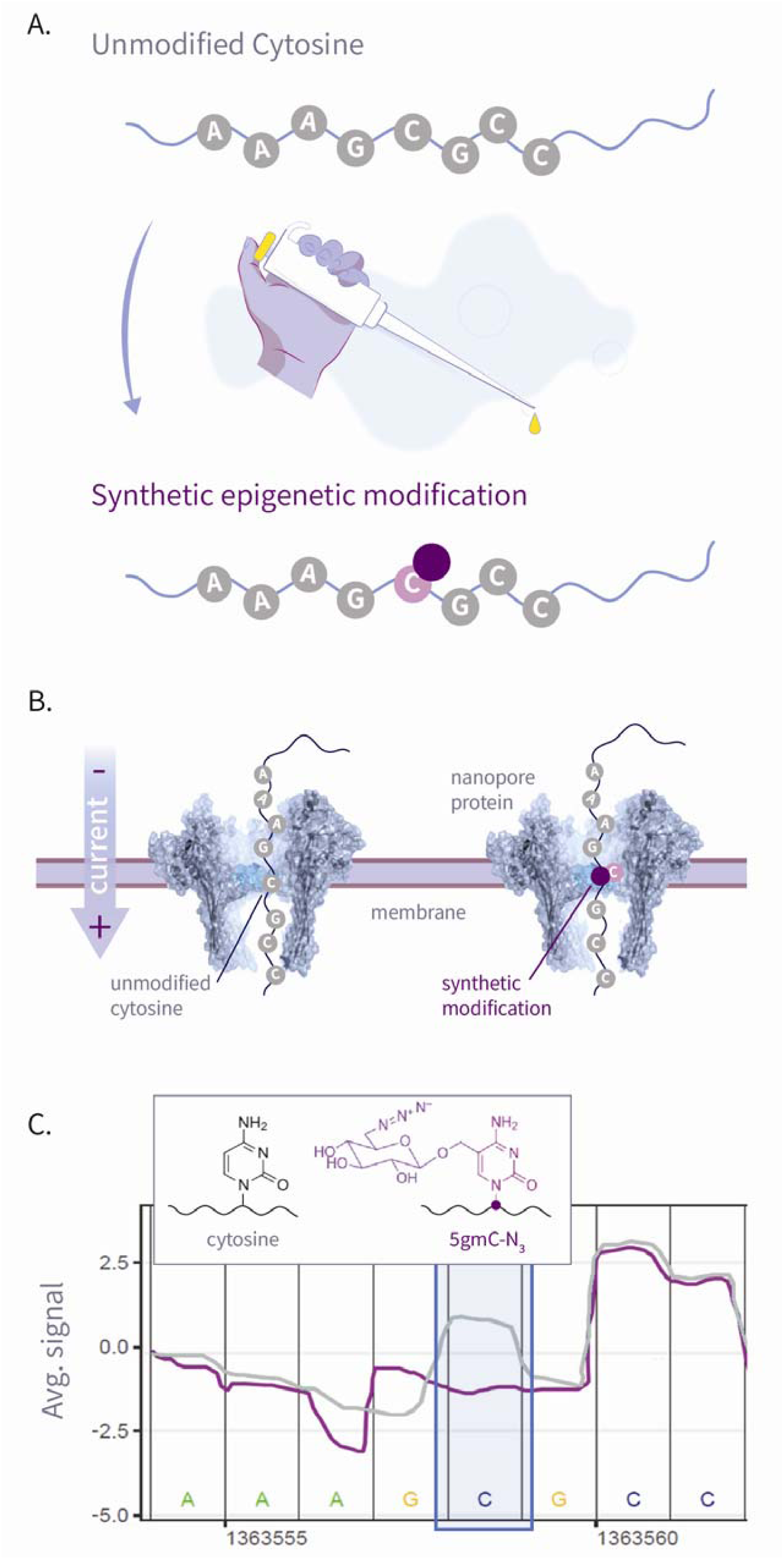
This plot shows an illustration of our write-and-read paradigm: (A) Write: chemoenzymatic conversion of designated genomic bases with synthetic epigenetic modifications. (B) Nanopore sequencing of the modified strands. (C) Read: detection of modified signal using modified-basecalling algorithms.

We first set out to explore DNA glucosylation as a contrast mechanism. Our study harnesses the natural glucosylation mechanism developed by bacteriophages^40^; we implemented an enzymatic glucosylation strategy using T4 β-glucosyltransferase (BGT) with either glucose or glucose-azide donors to selectively convert 5-hydroxymethylcytosine (5hmC) into either 5- glucosyl hydroxymethylcytosine (5gmC) or 5-azide-glucosyl-hydroxymethylcytosine (5gmC- N) ^38,41^. We demonstrate that these modifications elicit distinct ionic current signatures during nanopore sequencing, which are distinguishable from all three naturally occurring species of cytosine: unmodified cytosine, 5-methylcytosine (5mC), and 5-hydroxymethylcytosine (5hmC).

In addition, we demonstrate that other enzyme-substrate systems could be adapted to introduce a broad spectrum of chemical modifications at specific genomic sites. For example, methyltransferases, which catalyze the transfer of a methyl group to a nucleobase within their recognition sequences, could be repurposed for site-specific labeling^32^. Here, the natural methyl donor, S-adenosylmethionine (AdoMet), is replaced with a synthetic cofactor, 6-azido-S-adenosyl-L-methionine (6-N -AdoMet), which directly incorporates an azide moiety, facilitating downstream nanopore detection.

## RESULTS

### DNA GLUCOSYLATION DETECTION

The three common naturally occurring variants of cytosine, unmodified C, 5mC, and 5hmC, are the most studied via nanopore sequencing, with established models for multi-omic base calling^42^. To investigate our approach for chemoenzymatic labeling and detection of signal disturbances using nanopores, we initially examined the addition of two synthetic cytosine variants that can be chemoenzymatically generated, 5gmC and 5gmC-N (see Figure S1 for conversion schematics). In order to measure how the detected signal differs between the five cytosine variants, we amplified a 1kb portion of the λ-phage genome^43,44^ using PCR primers terminating with either C, 5mC or 5hmC. Then, sites bearing 5hmC sites were chemoenzymatically converted into 5gmC or 5gmC-N_3_, resulting in five amplicons carrying the same sequence but displaying the five cytosine variants for comparison (see Figure 2A). Finally, we barcoded each of the five amplicons and sequenced them on a single ONT MinION flowcell. Our chemoenzymatic tagging strategy, illustrated in Figure 2A, produced notable current shifts for 5gmC and 5gmC-N when compared to all natural variants of cytosine (C, 5mC, and 5hmC). Furthermore, the levels of 5hmC closely correspond with those of unmodified C, unlike the levels of 5mC, which is a previously documented phenomenon^45,46^.

**Figure 2.**
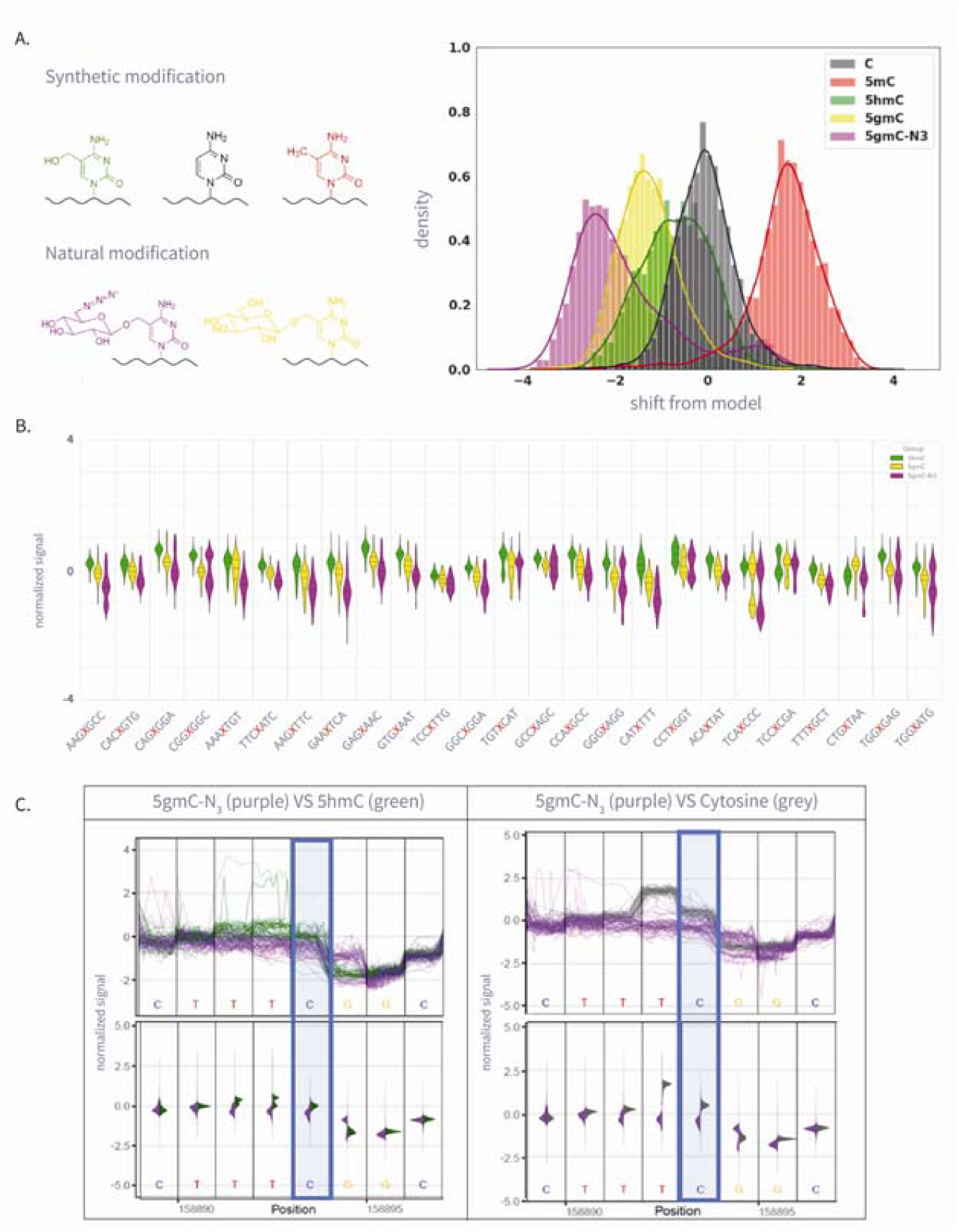
(A) The left panel is the molecular structure of various natural and synthetic cytosine derivatives: unmodified C, 5mC, 5hmC, 5gmC, and 5gmC-N□; The right panel is thesignal shift distributions of a single *k*-mer (GATGCG) from ONT’s trained model expected signal, which demonstrates clear differentiation between the natural and synthetic modifications. (B) The 25 Amplicon sequences showing normalized signal for 5hmC, 5gmC, 5gmC-N_3_. (C) Plots comparing ionic current signals for 5gmC-N□ (purple) versus 5hmC (green) on the left, and unmodified cytosine (grey) on the right. The y-axis represents normalized signal, and the x-axis denotes genomic position. Upper panels show raw signal traces, while lower density plots display signal distributions. The modified cytosine position of the CG context is highlighted in blue.

Following these initial results, we designed a panel of 25 amplicon sequences with varied sequence contexts flanking the target CpG site to test the effect of sequence context on the modified signal. For every sequence, we generated four PCR amplicons that differed only at the target CpG and contained the following nucleotides: C, 5hmC, 5gmC, and 5gmC-N . Then, we barcoded the samples and sequenced them on an ONT MinION flowcell.

Analysis of the ionic current signals revealed significant differences between 5gmC and 5gmC-N, as well as between these modified nucleotides and unmodified 5hmC (see Figure 2C). Since several bases occupy the pore at any given time, the measured signal reflects the integrated effect of the chain of nucleotides residing in the pore, known as the *k*-mer. A helicase releases the DNA through the pore base by base, and as the modified base changes its positions within the pore, the current shift changes. For simplicity, we focus on visualizing the eight positions around the modified cytosine. However, although signal variation can sometimes be detected up to eight positions away from the modification, the most profound disturbance is usually observed within two bases upstream of the modification site, and sometimes even within five positions upstream of it. This behavior, observed in both synthetic and natural modifications, aligns with previous findings regarding natural modifications^2^. The differences we detected were consistent across all pre-selected *k*-mers, as illustrated in Figure S2, and were verified via t-test statistics and effect size plots comparing the different samples (supporting information and Figures S3 and S4). Nevertheless, the exact shift patterns were highly sequence-specific, highlighting the need for extensive training before this approach can be implemented on a genome scale.

### DNA ALKYLATION DETECTION

To demonstrate the versatility of our write-and-read approach, we extended our experiments to include DNA alkylation reactions. Specifically, we used methyltransferases, which naturally alkylate specific sites on DNA by transferring methyl groups from the universal methyl donor S-adenosylmethionine (AdoMet) to particular nucleobases^30,47^. We employed both native and mutant methyltransferases, alongside synthetic cofactors, which facilitate the direct transfer of various chemical tags to DNA^30,39,48,49^; such tags could generate electrical contrasts useful for sequencing. For an expanded list of available AdoMet analogs, refer to Table S1. Furthermore, methyltransferases targeting non-canonical motifs, such as GpC or adenine, in mammalian genomes have been utilized to explore genomic information beyond traditional DNA genetics and epigenetics, including chromatin accessibility and protein binding^23,50,51^. We specifically focused on evaluating adenine alkylation, which could serve as a valuable marker for mapping protein-DNA interactions.

We generated PCR-amplified DNA products containing the TCGA motif, which serves as the recognition sequence for the M.TaqI methyltransferase that alkylates the terminal adenine^52^. We used M.TaqI along with AdoMet or an azide-functionalized analog, 6-N -AdoMet, to produce three variants of adenine in the TCGA-motif-containing DNA: unmodified, methylated (6mA), and azide-modified (N -A). For the reaction schematic, see Figure 3A. Next, the samples were barcoded and sequenced using a single ONT MinION flowcell.

**Figure 3.**
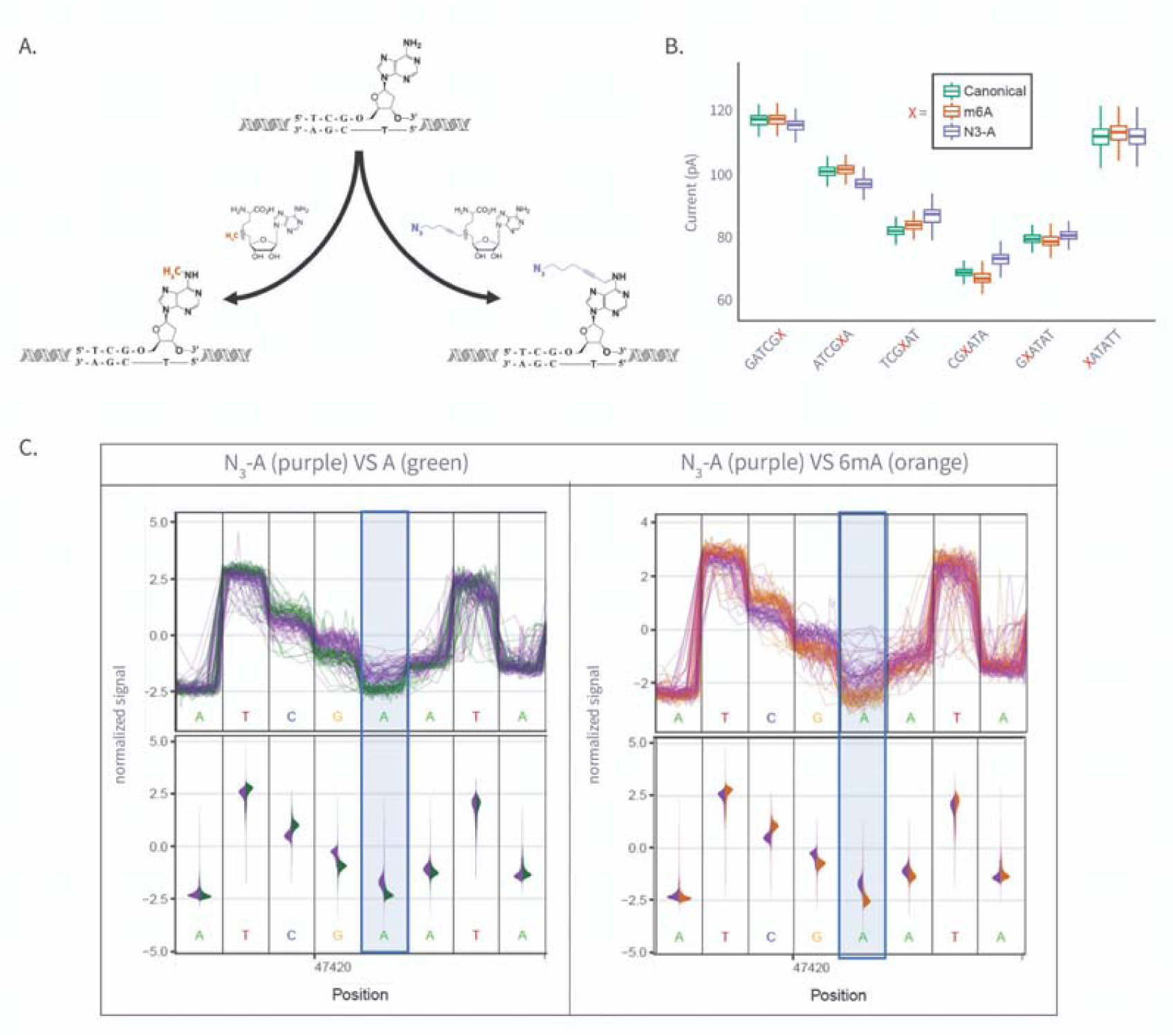
(A) Enzymatic transfer of the azide-methyl group to adenine by M.Taql methyltransferase using S-adenosylmethionine (AdoMet) co-factor and its azide-modified analog (Azide-AdoMet). (B) Box plots of ionic current (pA) for canonical adenine (*N* = 6569), N6-methyladenine (6mA, *N* = 7224), and azide-modified adenine (N□-A, *N* = 5191) across selected *k*-mers, showing distinct current shifts; A or modified-A position in the 5-mer is colored in red. (C) Plots comparing ionic current signals for azide-adenine (N□-A, purple) versus unmodified adenine (A, green; left) and N6-methyladenine (6mA, grey; right). The y-axis represents normalized ionic current (pA), and the x-axis denotes genomic position. Upper panels show raw signal traces, while lower violin plots display signal distributions. The modified adenine position is highlighted in blue.

Analysis of current distributions revealed distinct effects for 6mA and N -A (see Figures 3B and 3C; see Figure S5 for all modified locations). While the native A and 6mA overlap highly in signal in most positions, N -A exhibited more substantial shifts across multiple pore positions, as seen in both Figure 3B and 3C. This suggests that N -A is more easily detectable, likely due to its more considerable steric bulk and altered molecular interactions within the pore (for t-test Cohen’s d, see Figure S6).

In the absence of a model to call our modified bases, we further investigate the impact of our chemoenzymatic labeling on nanopore sequencing by examining events in the raw signal data that the signal realignment models could not confidently match with the reference genome. These unassigned events offer another perspective, highlighting instances where the observed ionic current significantly deviates from the expected patterns defined by the standard model. Since the model isn’t optimized for synthetic variants, these deviations suggest that these modifications cause measurable signal distortions. As shown in Figure S7 (glucosylation) and Figure S8 (alkylation), all our synthetic variants exhibit a higher rate of unassigned events compared to the native controls, up to seven bases upstream of the modified base. These findings suggest that synthetic modifications may cause signal disruptions distinct from those produced by naturally occurring nucleotides.

## DISCUSSION AND OUTLOOK

This study introduces an exogenous labeling method for nanopore sequencing based on the selective enzymatic manipulation of DNA with unnatural modifications. We demonstrate that the bases that we modify produce distinct electrical signatures as they translocate through the pore, broadening the palette of detectable electric signals. By extending the labeling toolbox we previously developed for fluorescent tagging of DNA^15,22,53^, we demonstrated the write-and-read concept using chemoenzymatic glucosylation or alkylation with an unnatural moiety that creates distinct electrical contrast. In the first case, we utilized β-glucosyltransferase (BGT) to selectively modify 5-hydroxymethylcytosine (5hmC) with glucose or engineered glucose-azide cofactors across diverse sequence contexts.

Similarly, our preliminary investigation into adenine modifications revealed that while both natural N6 methyladenine (6mA) and its azide-modified counterpart (N -A) produce measurable changes in ionic current, the azide modification consistently elicits a more pronounced effect in multiple sequence contexts. Although additional validation in diverse *k* mer environments is warranted, these results suggest that nanopore sequencing can accurately distinguish m6A and N -A as two orthogonal tags.

Despite the promising results, this study has several limitations. First, our experiments were primarily conducted on synthetic DNA substrates. While these provide controlled conditions for proof of concept validation, they may not fully recapitulate the complexity of native genomic DNA. Additionally, the increased occurrence of unassigned *k* mer events suggests that the chemical modifications, notably bulky adducts, can interfere with the basecalling accuracy of currently available models, complicating downstream analyses. A dedicated model for these unnatural modifications requires further optimization^46^ to ensure robust performance across various sequence contexts and modification densities. Furthermore, this project used the R9 MinION flow cells, which were discontinued in July 2024^54^. While these flow cells provided reliable data for the proof-of-concept, future work will re-establish the compatibility and performance of this chemoenzymatic labeling strategy with the newer-generation pore characteristics that may influence signal profiles^55,56^.

Protein pore engineering, electronics, and computational algorithms will continue to improve our ability to identify native modifications, such as DNA methylation. However, the presented approach offers the opportunity for signal engineering beyond the natural repertoire of nucleobases via selective chemical manipulation of DNA. Our future work will focus on creating training sets and robust computational models for calling synthetic modifications in different sequence contexts, similar to how current models are being developed to detect natural modifications accurately^46^. This will allow seamless integration of new signals beyond those naturally occurring on genomic DNA.

By expanding the molecular alphabet used in nanopore sequencing and incorporating synthetic modifications into recently developed methods for mapping chromatin structure^1,57^ and protein-DNA interactions^50,51,58^, we aim to enable the simultaneous recording of multiple genomic features in a single run. Previously, these techniques were limited by their dependence on canonical methylation cofactors, which constrained the variety of information obtainable from the same DNA molecule. However, as schematically illustrated in Figure 4, our approach could enable the concurrent analysis of the genetic sequence, DNA modifications, chromatin accessibility, and transcription factor binding sites, with binding sites labeled via antibody-mediated alkylation of proximal N_3_-A (using, for example, Dimelo^58^ or Champ^57^). Additionally, developing new chemistries for selective DNA modifications could open new avenues for studying the interactions among epigenetic marks, such as methylation, hydroxymethylation, and histone modifications, and their collective influence on gene regulation and cellular identity^59^. Therefore, our strategy provides a foundation for integrated multiomic analyses that capture genetic variation, dynamic epigenetic states, and chromatin organization in a single-pass readout, all while leveraging the benefits of long-read sequencing^42^.

**Figure 4.**
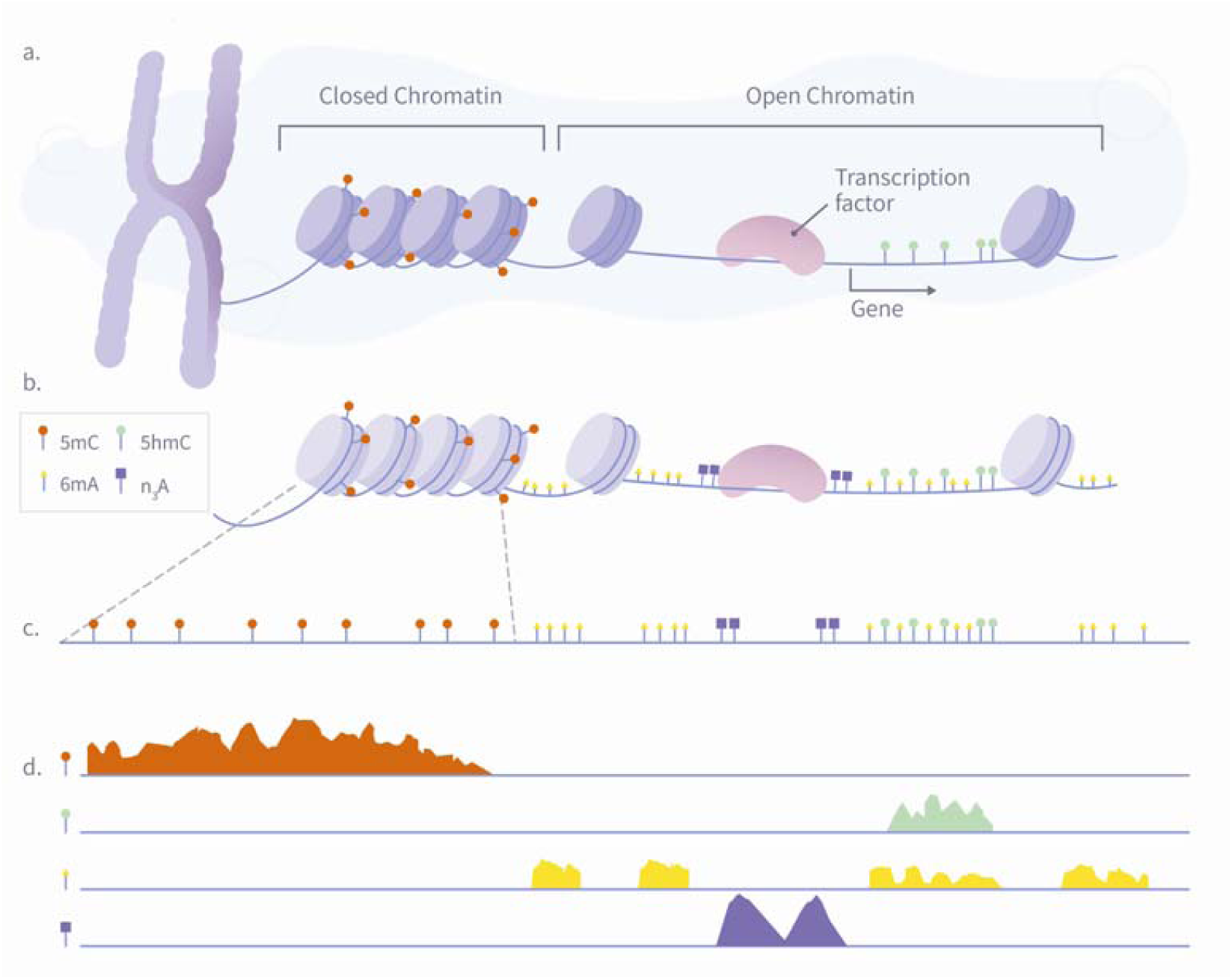
A schematic illustration of a single-pass multi-omic experiment demonstrating how nanopore sequencing can simultaneously detect multiple epigenetic and chromatin features. The diagram shows the process in four stages: (A) Nuclear chromatin is composed of genomic DNA, carrying native 5mC (red) and 5hmC (green) modifications, and associated with a plethora of DNA-binding proteins such as nucleosomes and transcription factors. (B) Exposed genomic DNA is labeled within permeabilized nuclei to mark protein footprints by utilizing the high frequency of Adenine, which provides high-resolution mapping. Transcription factor binding sites are marked with N_3_-A tags (purple) via antibody-mediated proximity labeling, while chromatin accessibility is marked by 6mA tags (yellow). (C) The DNA is extracted for nanopore sequencing, during which it is stripped from histones, transcription factors, and other bound proteins, maintaining both synthetic and natural modifications. (D) After sequencing and basecalling, the resulting data enable the simultaneous analysis of 5mC, 5hmC, chromatin accessibility, and transcription factor binding sites using standard bioinformatics tools.

## METHODS

### SAMPLES PREPARATION AND SEQUENCING

We began by performing PCR to generate site-specific cytosine variants across three λ- DNA fragments, as outlined in Table S2. Using 100 ng of template DNA and 0.4 µM of pre-ordered primers, which included unmodified cytosine, 5-methylcytosine (5mC), and 5- hydroxymethylcytosine (5hmC), we amplified products measuring 1.05 kb (10 003-11054), 2.80 kb (39608-42407), and 1.93 kb (46340-48266). The amplification was conducted using MyTaq™ Red Mix under the following program: 95 °C for 1 min; 30 cycles of 95 °C for 15 s, 58 °C for 15 s, and 72 °C for 2–3 min (extension time per fragment); final extension at 72 °C for 5 min; hold at 10 °C. Product size and concentration (26–37 ng/µL) were confirmed by agarose gel electrophoresis and Qubit fluorometry.

Purified amplicons were then enzymatically labeled (see Table S3 for reaction conditions). For 5hmC glycosylation of the 1.05 kb and 2.80 kb fragments, 30 µL reactions in NEBuffer 4 contained 1 µg DNA, 20U of T4 β-glucosyltransferase, and 45 µM UDP-glucose or UDP-6-N -glucose, incubated overnight at 37 °C. All reactions were purified via QIAquick PCR Purification Kit (Qiagen) spin columns and eluted in elution buffer. For adenine modifications on the 1.93 kb fragment, 75 µL reactions in NEB CutSmart buffer contained 0.26 mg/mL M.TaqI plus either 4.1 mM UDP-6-N -glucose or S-adenosylmethionine, incubated at 60 °C for 1 h, followed by proteinase K digestion (20 mg/mL) at 45 °C for 2h. The M.TaqI labeling reaction efficiency was assessed using agarose gel electrophoresis, as shown in Figure S9.

All the samples were barcoded using ONT’s EXP-NBD104 kit, end-repaired and adapter-ligated using the SQK-LSK109. Lastly, the prepared library was loaded onto ONT MinION FLO-MIN106 flow cells and sequenced using fast basecalling mode (See Table S4 for yield summary).

Then, to further validate and generalize our findings, 25 DNA fragments, each containing one of four classes of cytosine modifications, were prepared: C, 5hmC, 5gmC, and 5gmC-N . The genome of Escherichia coli K-12 MG1655^58^ was extracted using the Promega Wizard DNA Extraction Kit and used as a PCR template. Twenty-five unique 11-mer target sites were selected (Table S5), and forward primers were custom-synthesized to include a single 5hmC residue at the central position. PCR reactions were performed in 50 μL volumes, containing 100 ng of genomic DNA, 0.4 μM of each primer, 25 μL of Bioline MyTaq Red Mix (2×), and ultrapure water. Thermal cycling followed the program detailed in Table S6. Amplified products were purified with the QIAquick PCR Purification Kit (Qiagen) and checked for the correct size by agarose gel electrophoresis.

Selective glucosylation of 5hmC residues was performed in 1× NEBuffer 4 (New England Biolabs) using T4 β-glucosyltransferase. For standard labeling to produce 5gmC, 1 μg of purified PCR product was incubated with 20 units of enzyme and 45 μM UDP-glucose in a 30 μL volume at 37 °C overnight. For azide-functionalized labeling to produce 5gmC-N, the same conditions were used, but with 45 μM UDP-6-N -glucose as the cofactor; RP-HPLC confirmed the quantitative transfer of the azide-glucose moiety (Figure S10 and Table S7). After reaction, labeled DNA was purified using Zymo Oligo Clean & Concentrator columns and stored at -20 °C. These modified 11-mers were then barcoded using the EXP-NBD104 kit, library-prepared with the SQK-LSK109 kit, and sequenced on MinION FLO-MIN106 flow cells with fast basecalling (see Table S8 for a yield summary).

## DATA ANALYSIS

After sequencing, multi-read fast5 files were initially generated, each containing multiple reads. These files were converted into single-read fast5 format to facilitate downstream analysis using the "multi_to_single_fast5" utility from the "ont-fast5-API" library^59^. This conversion ensured that each fast5 file contained data from a single DNA molecule, as required for the subsequent signal analysis.

Basecalling was performed using ONT’s Guppy software (version 6.3.2) in high-accuracy mode (model dna_r9.4.1_450bps_hac), optimized for the R9.4.1 flow cell chemistry. Guppy translated the raw electrical signals into nucleotide sequences, generating FASTQ files (as shown in Table S4 and Table S8 yields’ summary).

To integrate the basecalled sequences with the raw ionic current data, we used Tombo’s annotate_raw_with_fastqs function^60^. This annotation step linked the FASTQ reads to the corresponding fast5 files, enabling a direct association between the nucleotide sequence and its underlying current profile, as required by Tombo’s 1.5.0 version.

Then we used Tombo’s resquiggle function, which first normalizes each read by subtracting its median (shift) and dividing by its median absolute deviation (scale), producing “normalized signal” levels. It then aligns the normalized signal (raw squiggle data) to the reference genome. After initial event detection and sequence assignment, Tombo refines the shift and scale parameters by fitting observed versus expected signal levels with a Theil–Sen slope for scale and the median of per base intercepts for shift. If either correction exceeds preset thresholds, the sequence to signal assignment is repeated until convergence. This per-read, sequence-dependent approach is more robust for samples with high modified-base content than mean-based methods and is now the recommended scaling procedure. Finally, the resquiggle process corrects for baseline drift and aligns the observed signals with the expected current values from the reference sequence, enabling precise mapping of signal shifts associated with the evaluated modifications. Alignments were performed against either the λ-phage^44^ or E.coli reference genome^58^.

The aligned, normalized, and resquiggled data was then analyzed to quantify the distinct current shifts induced by the chemical tagging. To visualize and compare the signal shifts across different modifications, we utilized Tombo’s plot command to generate plots showing the ionic current profiles for each modification type aligned along genomic positions. To conduct t-tests, assess the effect sizes, and extract descriptive statistics between group pairs, we used Tombo’s "detect_modifications" and "text_output" functions, respectively. To extract Tombo’s resquiggled normalized-signal level per genomic position, we used Tombo’s Python API.

To quantify and extract the raw signal and observed signal shifts, and to provide statistical validation of the chemical modifications, we utilized Nanopolish’s eventalign function.^2,61^. Nanopolish eventalign realigns the reads to the reference genome at the signal level, providing detailed information about the timing and magnitude of ionic current events at each base. The output from eventalign included precise event-level data for each read, such as the observed current mean, standard deviation, and dwell time, allowing for a comprehensive analysis of the modifications.

As seen in Figures S6 and S7, to further assess the impact of synthetic modifications on signal realignment, we examined the occurrence of unassigned *k*-mers, denoted as "NNNNNN", in the output of Nanopolish’s eventalign function. This function provides a position-by-position mapping of ionic current shifts, linking each detected signal to a specific nucleotide sequence. However, when the observed current deviates significantly from the expected signal profile, typically due to unmodeled DNA modifications, Nanopolish masks the *k*-mer as "NNNNNN", indicating that the raw signal could not be confidently aligned to the reference sequence. Importantly, this does not imply a failure in basecalling, as the ONT basecaller (e.g., Guppy) still assigns standard ACGT sequences; instead, it suggests that the signal perturbations introduced by the modifications led to uncertainty in realignment during downstream analysis.

## ASSOCIATED CONTENT

(1) Supplementary information accompanying this study, containing additional figures and tables referenced throughout the main text (PDF). (2) Figure S2 (PDF). (3) Figure S5 (PDF).

## AUTHOR INFORMATION

### CONFLICTS

WT has two patents (8,748,091 and 8,394,584) licensed to Oxford Nanopore Technologies. WT has received travel funds to speak at symposia organized by Oxford Nanopore. YE has a pending patent application concerning synthetic modifications for nanopore sequencing.

### ABBREVIATIONS

DNA: Deoxyribonucleic acid
RNA: Ribonucleic acid
PCR: Polymerase chain reaction
HPLC: High-performance liquid chromatography
ONT: Oxford Nanopore Technologies
BGT: β-Glucosyltransferase
Mtase: Methyltransferase
AdoMet: S-adenosylmethionine
SAM: S-adenosylmethionine
*K-*mer: oligonucleotide of length k
5hmC: 5-Hydroxymethylcytosine
5mC: 5-Methylcytosine
5gmC: 5-Glucosyl-methylcytosine
5gmC-N□: 5-Azide-glucosyl-methylcytosine
6mA: N□-Methyladenine
N□-A: Azide-modified adenine.

### Supporting Information Available

Supplementary figures (S1–S10), tables (S1–S8), additional methods, statistical details, raw nanopore current traces for cytosine and adenine modifications, analysis of non-assigned events, PCR and labeling conditions, and sequencing yield summaries.

