## Supplementary information file for "Write and Read: Harnessing Synthetic DNA Modifications for Nanopore Sequencing"

SUPPLEMENTARY MATERIAL

### Glucosylation of Cytosine

#### Figure S1

Enzymatic conversion of cytosine

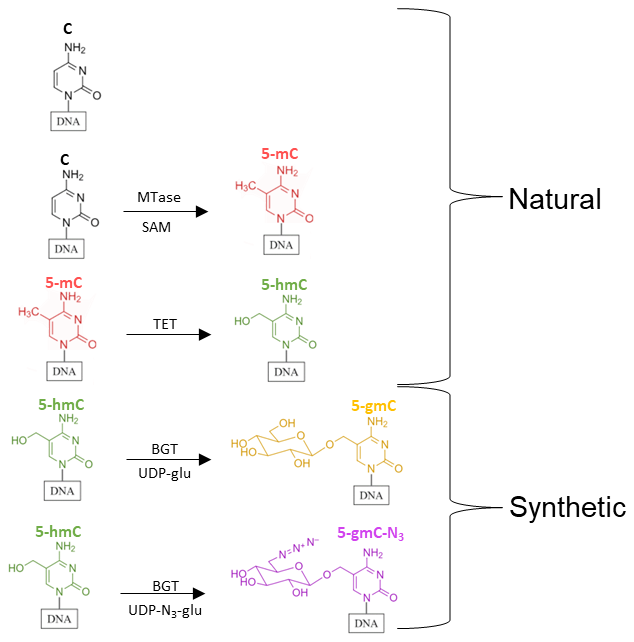

**Figure S1.** This schematic illustrates the conversion of a single DNA strand’s cytosine base at the C5 position, beginning with unmodified cytosine and progressing through natural epigenetic marks to synthetic labeling handles. First, a methyl group is added to form 5‑methylcytosine (5‑mC, red), which is then oxidized to 5‑hydroxymethylcytosine (5‑hmC, green). Then, either a glucose or glucose-ezide moiety is enzymatically attached to yield 5‑glyceryl‑methylcytosine (5‑gmC, gold), or 5‑gmC‑N₃ (purple).

#### Figure S2

This figure presents the complete set of raw nanopore current signals by Tombo’s plotting fuction (often referred to as “squiggle” traces) for unmodified cytosine (C), 5-hydroxymethylcytosine (5hmC), 5-glucosyl hydroxymethyl cytosine (5gmC), and 5-azide-glucosyl hydroxymethyl cytosine (5gmC–N₃). These data expand on the focused comparisons in the main text, where we discuss how glucosyl and azido-glucosyl groups on cytosine shift the ionic current measured by nanopore sequencing. By superimposing dozens of reads aligned to specific k-mers, Figure S1 illustrates that the modifications alter the shape and magnitude of the ionic current trace, yet these changes remain relatively consistent within a given modification type. As described in the Results section on DNA glucosylation, 5hmC retains current features somewhat like unmodified C, whereas 5gmC and 5gmC–N₃ cause more substantial and distinctive distortions. This raw-data overview helps confirm the reproducibility of the observed current differences and underscores the feasibility of detecting these synthetic modifications directly from the nanopore signal.

***Figure S2 is an attached PDF in the supplementary files due to figure dimensions.**

#### Figure S3 & S4

The D statistics were computed using the Tombo package's detect_modifications function with the level_sample_compare method. Tombo is a suite of tools developed by Oxford Nanopore Technologies for analyzing nanopore sequencing data. It is primarily used to identify modified nucleotides from raw nanopore sequencing reads. A Cohen’s D analysis quantifies the effect sizes of nanopore current shifts induced by various cytosine modifications. Unlike p-values, which only indicate whether a difference exists, Cohen’s D measures the magnitude of the difference relative to the variability in the data. Due to the high sequencing coverage per sample, the effect size metric was selected. Large Cohen’s D values support the conclusion that synthetic moieties—mainly the azido-glucose group—cause more pronounced and consistent deviations in ionic current.

D statistic distribution for 5gmC-N3 t-tests.
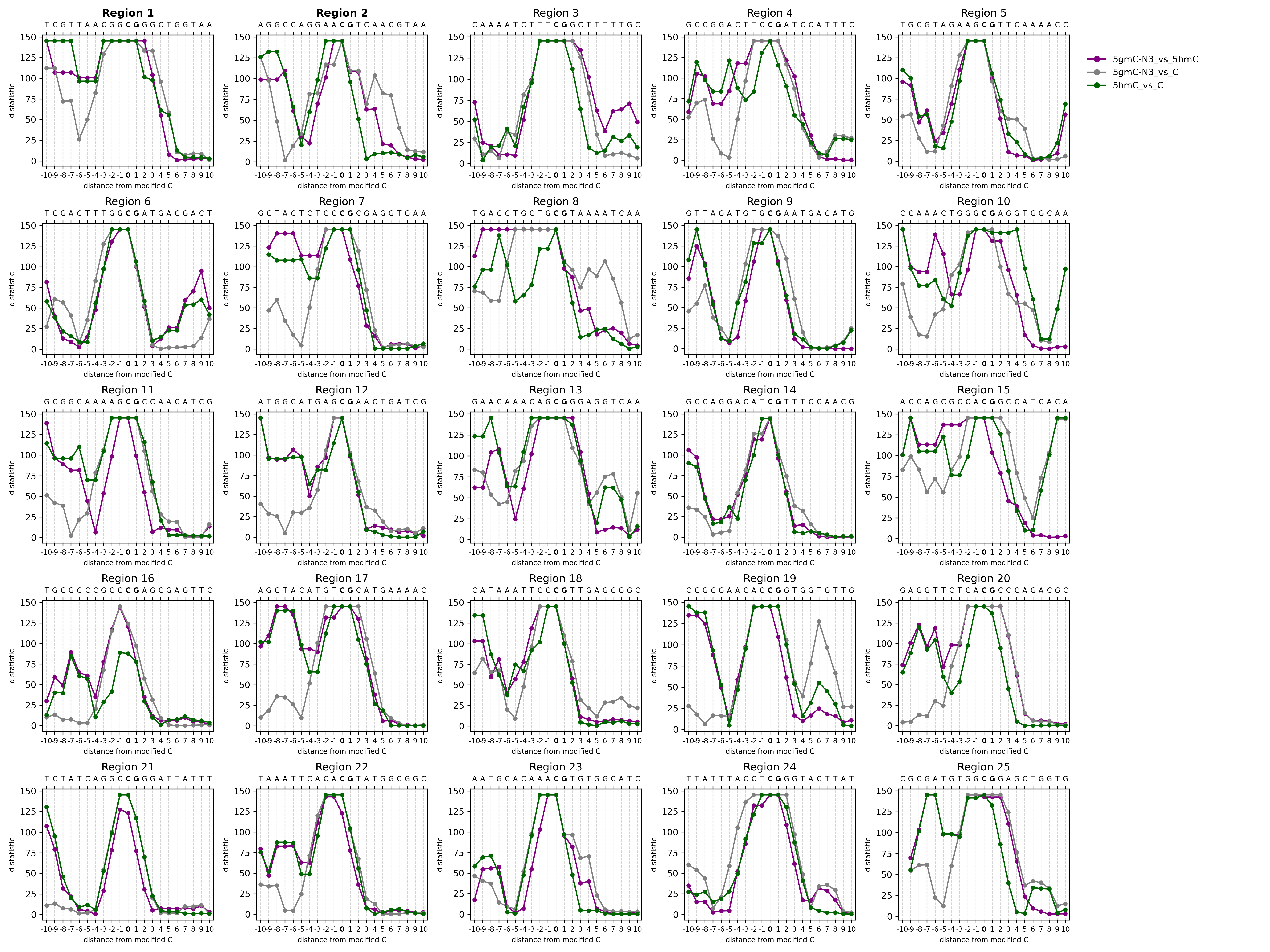

**Figure S3.** The panel plot displays the D statistic values for 25 modified genomic regions, with each subplot representing a distinct region with the modified kmer. The y-axis shows the D statistic of a t-test between groups, while the x-axis presents the nucleotide sequence (top) and the relative distance from the central position (bottom), ranging from –10 to +10 bases; CG context marked in bold. The presented groups are: 5gmC-N3 reads vs. 5hmC reads in purple; 5gmC-N3 reads vs. C reads in gray; 5hmC reads and C reads, for a natural-modification reference, in green. The Tombo package has a D statistic upper limit of 145.2219.

D statistic distribution for 5gmC t-tests.

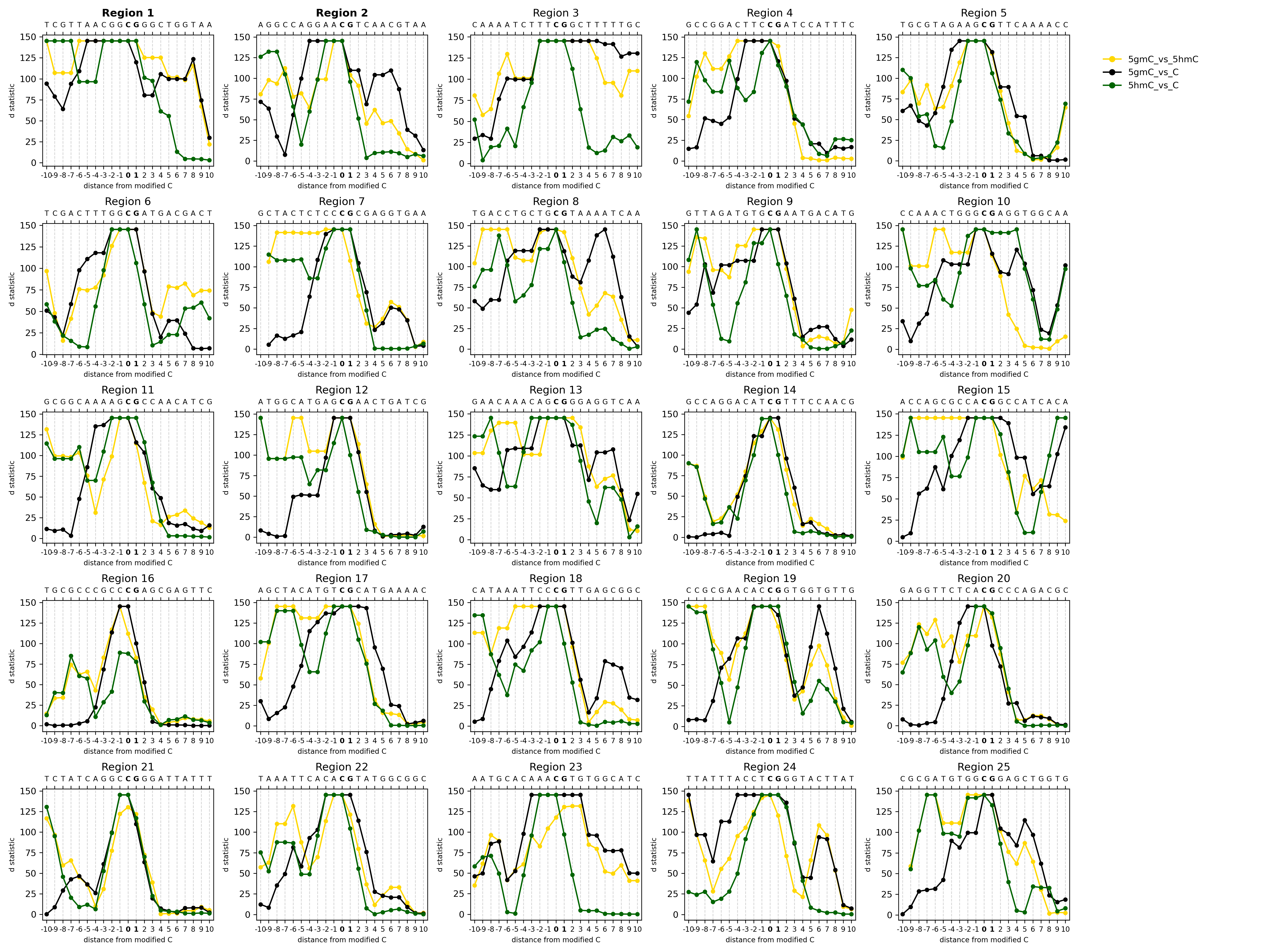

**Figure S4.** Similar to figure S3, but for 5gmC t-tests. The presented groups are: 5gmC reads vs. 5hmC reads in yellow; 5gmC reads vs. C reads in black; 5hmC reads and C reads in green for a natural-modification reference.

### Alkylation of Adenine

#### Table S1

In our “write-and-read” strategy for nanopore sequencing, methyltransferases play a key role by transferring methyl groups, or alternative functional moieties, from S-adenosylmethionine (AdoMet) to DNA. By substituting natural AdoMet with synthetic analogs, it becomes possible to incorporate diverse chemical tags, such as azides or alkynes, directly into specific genomic sites. This expands the range of detectable DNA modifications beyond native methylation, enabling the generation of distinct ionic signatures in nanopore reads. This table, which is a kind donation from Prof. Elmar Weinhold, summarizes several AdoMet analogs that vary in their substituents and heteroatoms, thereby providing different reactive handles or steric properties. These analogs are central to our approach because they allow precise, sequence-specific “writing” of unnatural modifications onto DNA. As demonstrated in this work, such modifications can be “read” through nanopore sequencing, yielding characteristic electrical shifts and paving the way for multi-omic sequencing.

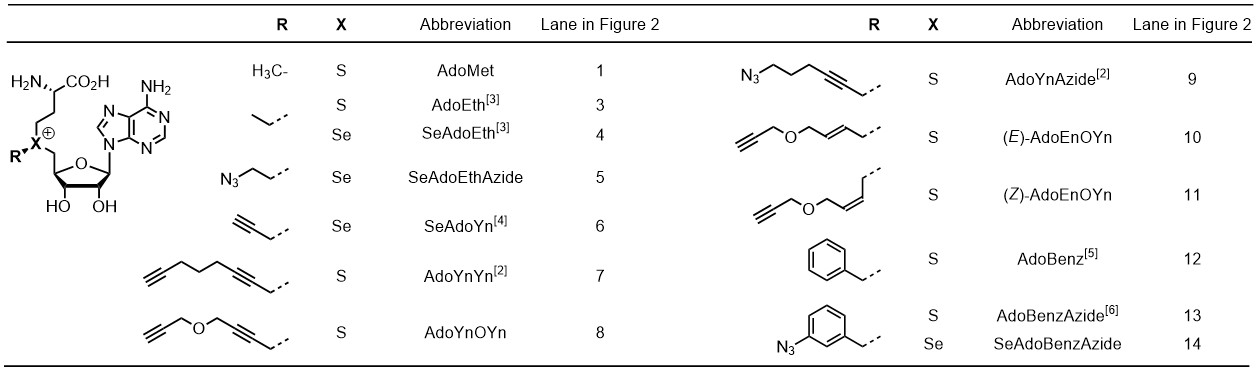

#### Figure S5

Similar to figure S1, this figure presents raw nanopore current signals generated by Tombo’s plotting fuction (“squiggle” plots) for unmodified adenine (A), N6-methyladenine (6mA), and azide-adenine (N₃-A). Reference genome used is of Enterobacteria phage lambda (NC_001416.1).

***Figure S5 is an attached PDF in the supplementary files due to figure dimensions.**

#### Figure S6

D statistic distribution for 5gmC-N3 t-tests.

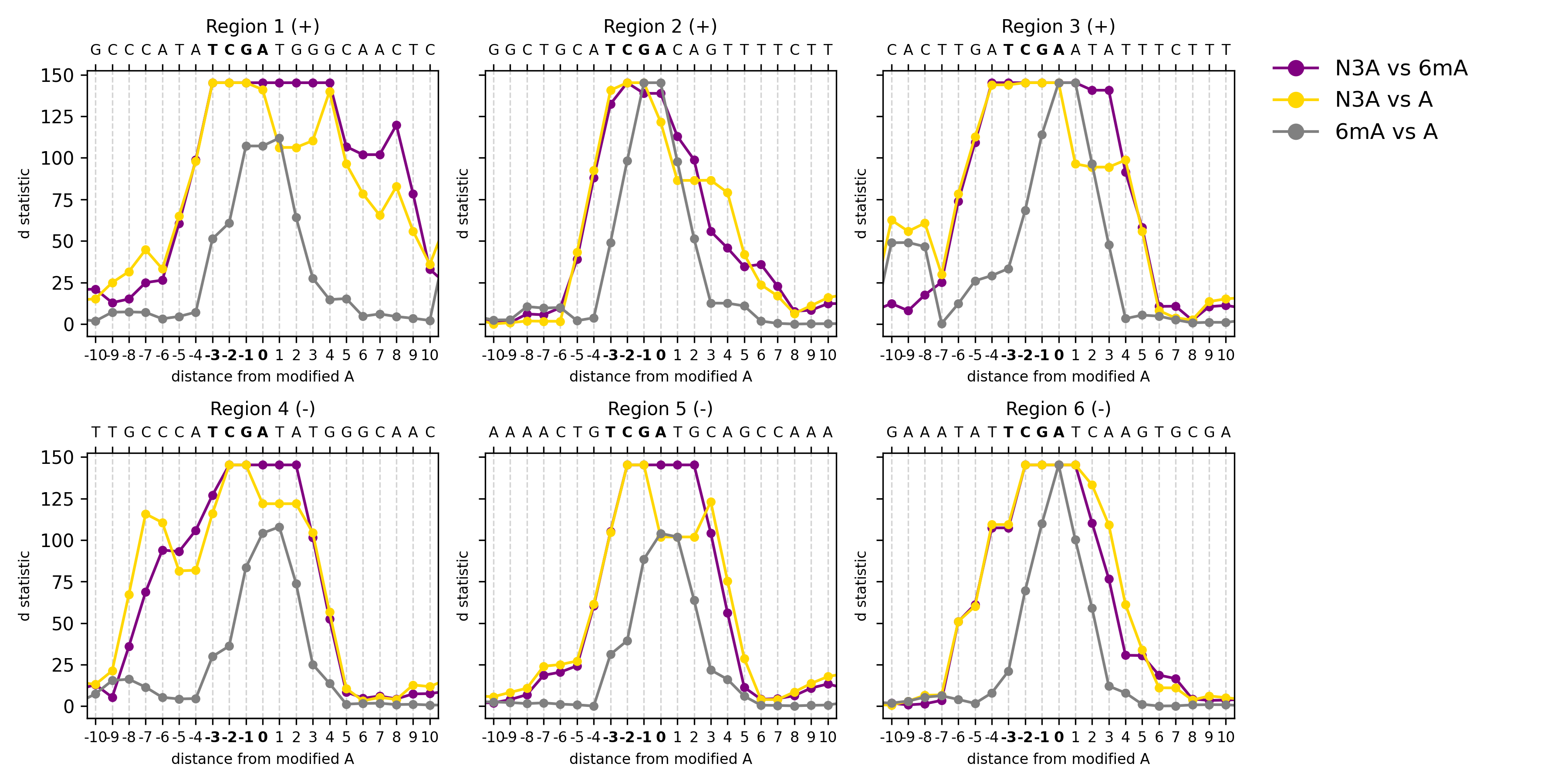

**Figure S6.** This plot displays the D statistic values for three modified genomic regions. Similar to Fig S2 & S3, the y-axis shows the D statistic of a t-test between groups, while the x-axis presents the nucleotide sequence (top) and the relative distance from the central position (bottom), ranging from –10 to +10 bases; M.Taq’s TCGA motif is marked in bold. The presented groups are: N3-A reads vs. 6mA reads in purple; N3-A reads vs. A reads in yellow; 6mA reads and A reads, for a natural-modification reference, in gray. The Tombo package has a D statistic upper limit of 145.2219.

### Analysis of Non-Assigned Events

#### Figure S7

The main text notes that bulky additions to 5hmC (i.e., 5gmC–N₃) often generate signals that deviate from the canonical basecaller model. To further examine these effects, we analyzed basecalling errors using Nanopolish, a widely used tool for signal-level analysis of nanopore sequencing data. One of its key functions, eventalign, maps each observed current signal to a specific k-mer in the reference sequence, providing a base-resolved view of how modifications affect the nanopore signal. In cases where the observed ionic current does not match any expected k-mer in the trained model, Nanopolish eventalign assigns the sequence “NNNNNN” to indicate an unrecognized event. Here, we plotted the frequency of these “N” events for each modification type (C, 5hmC, 5gmC, and 5gmC–N₃). Higher “NNNNNN” rates in sequences containing 5gmC–N₃ suggest that this synthetic modification introduces a distinct and unpredictable electrical signature, disrupting standard basecalling. This metric provides evidence that nanopore sequencing can distinguish chemically modified bases, as synthetic tags significantly alter the expected current profiles, making them harder for existing basecallers to interpret.

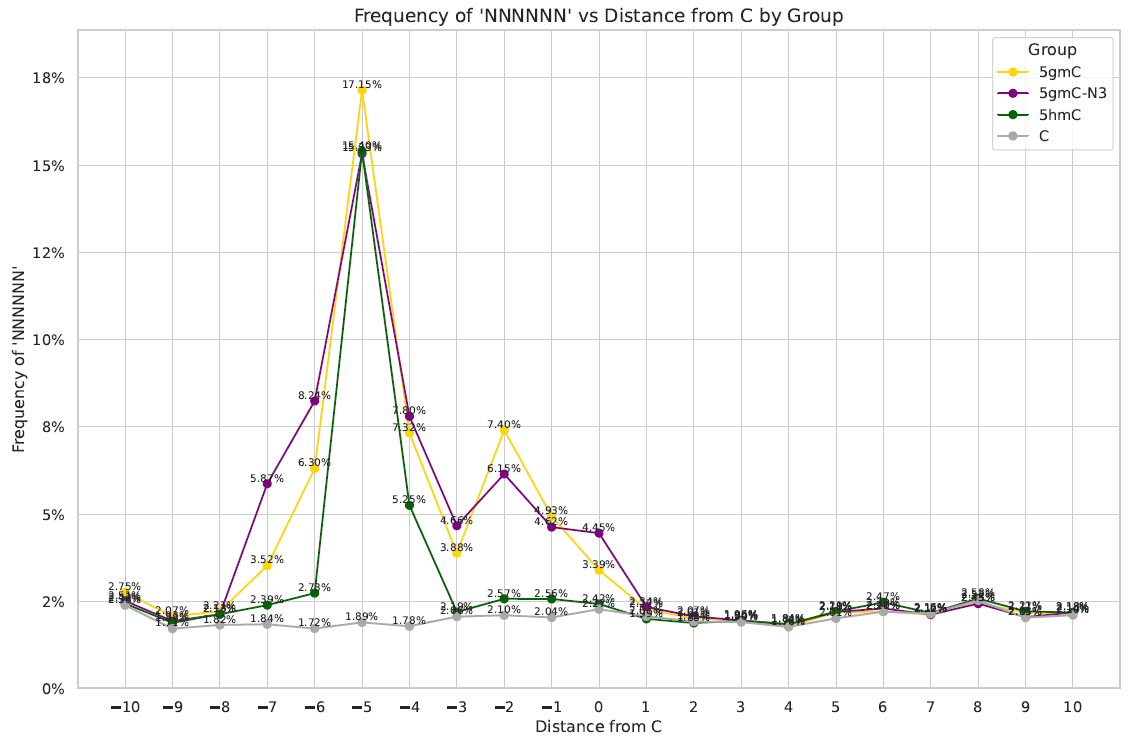

**Figure S7.** Frequency of "NNNNNN" k-mers in nanopore basecalling across different modification states. The x-axis represents the distance from the modified site (C/5hmc/5gmc/5gmC-N3), while the y-axis shows the percentage of total events labeled as "NNNNNN".

#### Figure S8

Here we plotted the frequency of “N” events for each Adenine modification type (N₃-A, 6mA. A). As in figure S4, higher “NNNNNN” rates in sequences containing suggest that this synthetic modification introduces a distinct electrical signature. This metric provides additional evidence that nanopore sequencing can distinguish chemically modified bases, as synthetic tags significantly alter the expected current profiles, making them harder for existing basecallers to interpret.

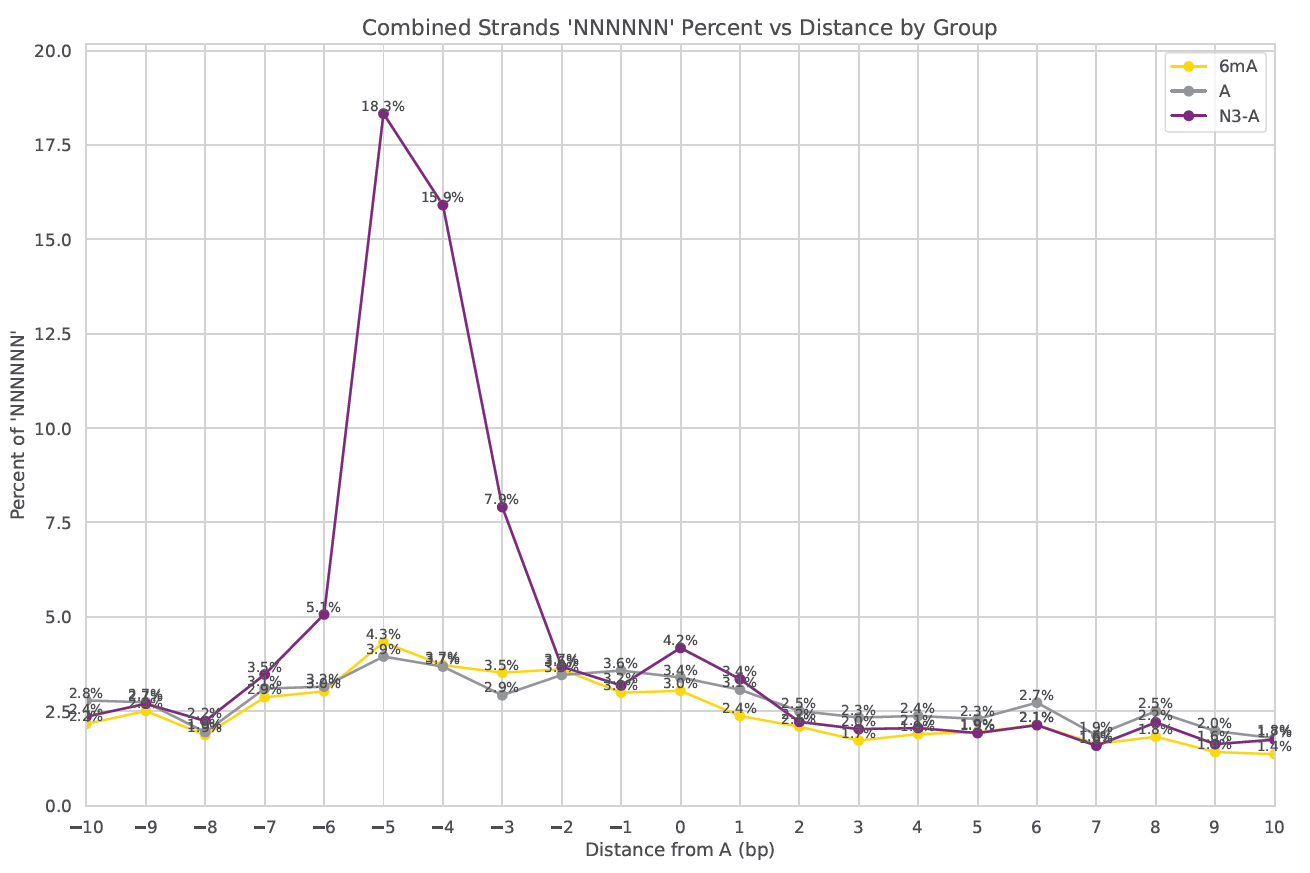

**Figure S8.** Frequency of "NNNNNN" k-mers in nanopore basecalling across different modification states. The x-axis represents the distance from the modified site (A/6mA/N3-A), while the y-axis shows the percentage of total events labeled as "NNNNNN".

### Methods

#### Table S2

A summary of the PCR conditions used to generate site‐specific cytosine, 5-methylcytosine (mC), and 5-hydroxymethylcytosine (5-hmC) variants in three λ‐DNA fragments.

**Table S2.**

Primer pairs and PCR product sizes for λ‐DNA fragments

| # | Start | End | Primer F (5′→3′) | Primer R (5′→3′) | Tₘ (°C) | PCR product (bp) |
| --- | --- | --- | --- | --- | --- | --- |
| 1 | 39608 | 42407 | TAACTTGACCGTGAGCAGATGCGTC | ATCATCTTCTTCCTCGTGCATCGAGC | 60 | 2 799 |
| 2 | 10003 | 11054 | CTCATGCTGAAAACGTGGT | GGACAGGACCAGCATACG | 53 | 1 051 |
| 3 | 46340 | 48266 | CTCAGAGAGAGGCTGATCACTA | GAACAACAACCCGCAACATC | 55 | 1 926 |

***Note.*** All reactions were run on a Bio-Rad C1000 with the following program: initial denaturation at 95 °C for 1 min; 30 cycles of 95 °C for 15 s, 58 °C for 15 s, and 72 °C for the time indicated in Table 1; final extension at 72 °C for 5 min; hold at 10 °C.

#### Table S3

To distinguish and track specific base modifications, we employed two enzymatic labeling strategies on the purified λ-DNA amplicons. For 5-hydroxymethylcytosine in the 1 051 bp and 2 799 bp fragments, T4 β-glucosyltransferase was used to transfer either natural glucose or azide-tagged glucose onto the 5-hmC residue. For adenine at TCGA motifs (fragment 1 926 bp), M.TaqI methyltransferase was used either with S-adenosylmethionine to install a methyl group or with UDP-6-N₃-glucose to introduce an azide-functionalized glucose.

**Table S3.**

Enzymatic labeling reaction conditions

| **Application** | **Fragment (bp)** | **Volume (µL)** | **Enzyme (amount)** | **Cofactor (conc)** | **DNA input** | **Incubation** |
| --- | --- | --- | --- | --- | --- | --- |
| 5-hmC glycosylation  (5-gmC) | 1051 &  2799 | 30 | T4  β-glucosyltransferase (20 U) | UDP-glucose  (45 µM) | 1 µg | 37 °C, overnight |
| 5-hmC glycosylation  (5-gmC-N₃) | 1051 &  2799 | 30 | T4  β-glucosyltransferase (20 U) | UDP-6-N₃-glucose  (45 µM) | 1 µg | 37 °C, overnight |
| Adenine labeling  (methyl) | 1926 | 75 | M.TaqI  (0.26 mg/mL) | S-adenosylmethionine | 4.37 µL (343 ng/µL) | 60 °C, 1 h;  then proteinase K  (20 mg/mL), 45 °C, 2 h |
| Adenine labeling  (azide) | 1926 | 75 | M.TaqI  (0.26 mg/mL) | UDP-6-N₃-glucose  (4.1 mM) | 4.37 µL (343 ng/µL) | 60 °C, 1 h;  then proteinase K  (20 mg/mL), 45 °C, 2 h |

***Note.*** All reactions were purified via QIAquick PCR Purification Kit and eluted in EB buffer before downstream processing.

#### Table S4

This table describes the yields of the nanopore sequencing similar to Table S3. In this sequencing features the adenine data presented in this paper. The experiment used barcoding where different regions where labeled with different modifications.

**Table S4.**

Yields of the nanopore sequencing for each sample.

| Barcode | Sample | GB | Passed reads | Failed reads | Percent failed reads |
| --- | --- | --- | --- | --- | --- |
| 01 | Unmodified | 0.069 | 35729 | 1134 | 3.08% |
| 02 | 5mC/6mA | 0.085 | 50443 | 1634 | 3.14% |
| 03 | 5hmC | 0.069 | 36690 | 1062 | 2.81% |
| 04 | 5gmC | 0.080 | 44062 | 1392 | 3.06% |
| 05 | 5gmC-N3/N3-A | 0.084 | 46202 | 1447 | 3.04% |

***Note.*** The “GB” column indicates each sample's total sequencing data generated (in gigabytes). “Passed reads” denotes the number of successfully basecalled reads, while “Failed reads” represents the reads that did not meet the quality thresholds for accurate basecalling. The “Percent failed reads” is calculated as the proportion of failed reads relative to the total number of reads (passed + failed).

#### Table S5

The following table provides a list of the 25 selected 11-mer sequences and their genomic coordinates, which served as the target regions for our chemoenzymatic glucosylation-based labeling and nanopore sequencing experiments.

**Table S5.**

Summary of 25 unique 11-mer target sites.

| # | Start (C) | End (G) | 11-mer | Primer F | Primer R | TM | PCR product (bp) |
| --- | --- | --- | --- | --- | --- | --- | --- |
| 1 | 101223 | 101224 | AACGGCGGGCT | CTTCGTTAACGGCGGGCTG | ATACGCGTCAATACCGCCTT | 61 | 1136 |
| 2 | 148465 | 148466 | AGGAACGTCAA | GAAAGGCCAGGAACGTCAAC | GGCTTAACGTCATGCTGGTG | 62 | 1150 |
| 3 | 158893 | 158894 | TCTTTCGGCTT | GGTTACGTCAAAATCTTTCGGCTT | AGAAGATATCGACCCGCCTC | 58 | 1077 |
| 4 | 189720 | 189721 | ACTTCCGATCC | CGCCGGACTTCCGATCCATT | TGTGATCTGCCCGGCATTAAA | 61 | 1000 |
| 5 | 192927 | 192928 | AGAAGCGTTCA | CAAATGCGTAGAAGCGTTCAAA | CGCAGGTAACCAGTGATTCT | 62 | 1002 |
| 6 | 337959 | 337960 | TTTGGCGATGA | ACCTTCCTCGACTTTGGCGATGAC | TCGCCGATAATTTTCATCGCGTC | 64 | 1054 |
| 7 | 628537 | 628538 | TCTCCCGCGAG | GCTACTCTCCCGCGAGGTGAAATAA | GTGTGTTGCATTAAGTTCTGTGCC | 63 | 1040 |
| 8 | 639392 | 639393 | TGCTGCGTAAA | GTCTGACCTGCTGCGTAAAA | CCAAAACGTTCGCCCATC | 62 | 1014 |
| 9 | 798064 | 798065 | ATGTGCGAATG | CCCAGTTAGATGTGCGAATGA | CAGACATCCACTGAGCTGTTTA | 62 | 1144 |
| 10 | 1329610 | 1329611 | CTGGGCGAGGT | AAGCCAAACTGGGCGAGGT | CAGTAAGGCGCTCTTGCCA | 62 | 1042 |
| 11 | 1363558 | 1363559 | AAAAGCGCCAA | GGGCGGCAAAAGCGCCAA | TATCATTCATCGCCACAAACAGGC | 63 | 1080 |
| 12 | 1457502 | 1457503 | ATGAGCGAACT | GATGGCATGAGCGAACTGAT | CCAGTGGGTTATAGAAACGTTCA | 62 | 1047 |
| 13 | 1674898 | 1674899 | AACAGCGGGAG | CTTGAACAAACAGCGGGAGG | ACCATTGATGCTGCCAAACC | 60 | 1018 |
| 14 | 1863202 | 1863203 | GACATCGTTTC | GGCCAGGACATCGTTTCCAA | GTCGTGCGGCCAGAATTTTT | 62 | 1086 |
| 15 | 2000337 | 2000338 | CGCCACGGCCA | AACCAGCGCCACGGCCAT | AATGTTGCTCGACACCAATAACAG | 63 | 1063 |
| 16 | 2031744 | 2031745 | CCGCCCGAGCG | TGCGCCCGCCCGAGCGA | TTGTGAGCCTGGCGAAAAATCATC | 63 | 1164 |
| 17 | 2201063 | 2201064 | CATGTCGCATG | CCAGCTACATGTCGCATGAA | CCAGTTCTGTTGCGCCAAAA | 60 | 1167 |
| 18 | 2238240 | 2238241 | ATTCCCGTTGA | GGCATAAATTCCCGTTGAGC | CTTGTTGGAAATGAGCGGTATC | 62 | 1027 |
| 19 | 2337263 | 2337264 | AACACCGGTGG | AATACCGCGAACACCGGTGG | CCGTCGTACTATTTTCGAACTGC | 63 | 1084 |
| 20 | 2659799 | 2659800 | TCTCACGCCCA | CCGGAGGTTCTCACGCCCAG | GACCTGATGTCTTTCTCCGGTCAC | 63 | 1155 |
| 21 | 2830766 | 2830767 | CAGGCCGGGAT | GATTTCTATCAGGCCGGGATTA | TGCTGATTGAACAACTGGAAAG | 62 | 1057 |
| 22 | 2881691 | 2881692 | TCACACGTATG | GCTGTAAATTCACACGTATGGC | AATAAGCACATTAGCGCTTGC | 62 | 1038 |
| 23 | 2904823 | 2904824 | ACAAACGTGTG | GGTTTCAATGCACAAACGTGTG | GGCAATTTGCACTTGGGTAT | 59 | 1084 |
| 24 | 3274919 | 3274920 | TACCTCGGGTA | CCAGAATCATTTTATTTACCTCGGGTACT | CGATCTGCAAAATATCCTCGACCA | 61 | 1168 |
| 25 | 3303866 | 3303867 | TGTGGCGGAGC | CGCGATGTGGCGGAGCTG | GCTGGGTTTGTTTGACTATCTGGC | 63 | 1057 |

***Note.*** The "Start (C)" and "End (G)" columns indicate the genomic coordinates corresponding to the first (cytosine) and the last (guanine) nucleotide of each 11-mer, respectively. The "11-mer" column lists the exact nucleotide sequences selected for analysis. The "Primer F" and "Primer R" columns list the forward and reverse primers used for amplification. The "TM" column indicates the melting temperature of each primer pair, and the "PCR product (bp)" column shows the size of the PCR product in base pairs.

#### Table S6

A detailed breakdown of the thermal cycling steps used to amplify DNA fragments harboring 5hmC for subsequent glucosylation and azido-glucosylation is included here. The table lists each temperature, duration, and cycle count for generating the target PCR amplicons, as described in the Methods. Careful temperature control and cycle numbers ensure efficient amplification while preserving 5hmC integrity for later enzymatic labeling to form 5gmC and 5gmC–N₃.

**Table S6.**

PCR steps for preparation of DNA fragments with synthetically introduced modifications

| Step | Temperature (°C) | Time (min) |
| --- | --- | --- |
| 1 | 95 | 1:00 |
| 2 | 95 | 0:15 |
| 3 | 58 | 0:15 |
| 4 | 72 | 3:00 |
| 5 | Repeat steps 2-4 (x30) | - |
| 6 | 72 | 5:00 |
| 7 | 10 | Pause |

#### M.TaqI Labeling Reaction

We conducted a restriction-protection assay to evaluate the efficiency of M.TaqI in transferring the azide cofactor AdoYnAzide onto PCR fragments and to identify the lowest cofactor concentration that still provides complete protection. In each reaction, 500 ng of PCR product was incubated with a fixed amount of M.TaqI and serial twofold dilutions of AdoYnAzide (1:1, 1:2, 1:4, 1:8, 1:16, 1:32, 1:64, 1:128, and 1:256 enzyme-to-site ratios). Labeling was performed at 65°C for 1 hour, followed by a 1-hour Proteinase K treatment at 45°C to inactivate the enzymes. All samples were then digested with R.TaqI, which cleaves only unmodified recognition sites. Before loading, a non-SDS loading dye was added, and the reactions were separated on a 1% agarose gel (95 V, 80 min). Gels were subsequently stained with SYBR Safe. Fully labeled DNA appears as the intact 1,217 bp band, while incomplete labeling results in a 500 bp cleavage product. Complete protection was observed at the 1:8 dilution, with protection decreasing at higher dilutions, indicating that M.TaqI-mediated transfer of AdoYnAzide is effective down to approximately 1:8 but reduces at lower cofactor concentrations.

##### *Figure S9*

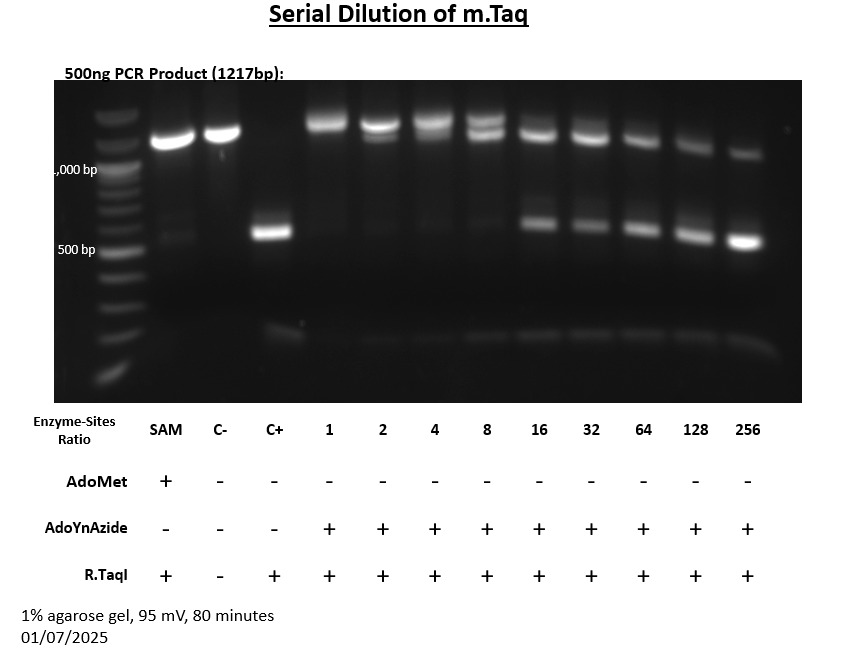

**Figure S8**. Azide-labeling protection assay on a 1,217 bp PCR fragment.

#### 5hmC labeling Efficiency

5hmC residues were labeled via a chemoenzymatic reaction. In each reaction tube, 1 μg of genomic DNA was mixed with 3 μL of 10X buffer 4 (New England Biolabs), uridine diphosphate-6-azide-glucose (UDP-6-azidoglucose) at a final concentration of 45 μM, 2 μL (20 units) of T4 phage β‑glucosyltransferase (T4‑βGT, New England Biolabs), and ultrapure water to bring the final volume to 30 μL. The reaction mixture was incubated overnight at 37 °C. The labeled DNA was then purified from excess reagents using Oligo Clean & Concentrator columns (Zymo Research) according to the manufacturer’s protocol, with three washing steps and two elutions for optimal recovery. For best yield, no more than two micrograms of DNA (from two combined reaction tubes) were loaded per column, and samples were stored at 4 °C until further analysis.

##### *Figure S10*

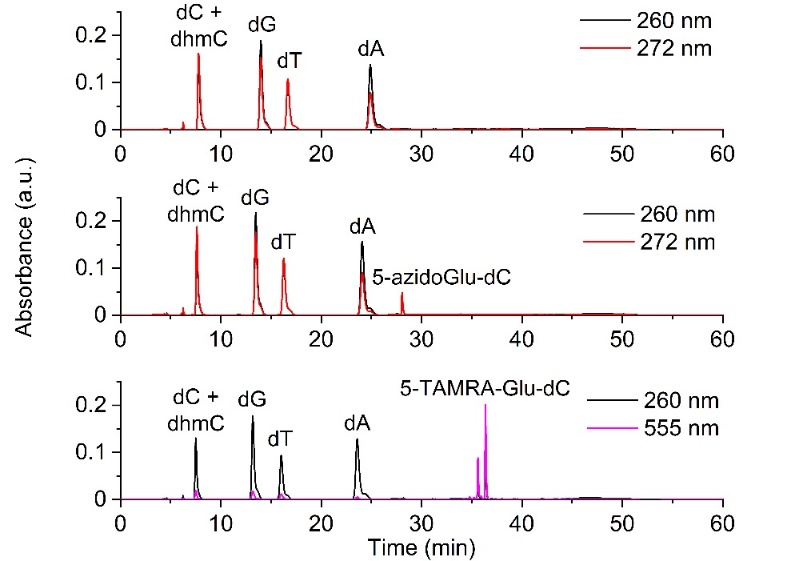

**Figure S8.** 5hmC labeling analysis by RP-HPLC of nucleosides obtained after enzymatic fragmentation of duplex ODN^7^. X axis is Time (min); Y axis is absorbance (a.u.). Top: Control reaction with duplex ODN containing one 5hmC residue. Bottom: Modification of the duplex ODN with T4-βGT and UDP-6-azidoglucose.

##### *Table S7*

**Table S7.** Quantification of 5hmC labeling analysis by RP-HPLC analysis^7^. Amounts of nucleosides obtained after duplex modification with T4-βGT and UDP-6-azidoglucose.

|  | dC +  5hmdC | dG | dT | dA | 5-azido-Glu-dC | Yield [%] |
| --- | --- | --- | --- | --- | --- | --- |
| Control | 7.0 | 7.0 | 6.0 | 6.0 | - |  |
| 5gmC-N3 | 6.0 | 7.0 | 6.3 | 5.8 | 1.1 | 100 |

#### Table S8

This table describes the yields of the nanopore sequencing for each sample type used in our chemoenzymatic labeling experiments. The table lists the data volume (in gigabytes), the number of reads that passed guppy basecalling, the number of reads that failed basecalling, and the corresponding percentage of failed reads. These metrics help illustrate how the different modifications affect the nanopore basecalling process. Notably, modifications tend to have a higher percentage of failed reads, reflecting the additional signal perturbations introduced by these modifications.

**Table S8.**

Yields of the nanopore sequencing for each sample.

| Sample | GB | Passed reads | Failed reads | Percent failed reads |
| --- | --- | --- | --- | --- |
| Cytosine | 0.37 | 309651 | 2090 | 0.60% |
| 5hmC | 0.25 | 211495 | 14284 | 6.75% |
| 5gmC | 0.35 | 295982 | 20283 | 6.85% |
| 5gmC-N3 | 0.25 | 212337 | 15375 | 7.24% |

***Note.*** The “GB” column indicates each sample's total sequencing data generated (in gigabytes). “Passed reads” denotes the number of successfully basecalled reads, while “Failed reads” represents the reads that did not meet the quality thresholds for accurate basecalling. The “Percent failed reads” is calculated as the proportion of failed reads relative to the total number of reads (passed + failed).
