## Supplementary material for "Write and Read: Harnessing Synthetic DNA Modifications for Nanopore Sequencing": Figure_S5

| Modified A Position (NC_001416.1) | Sequence | Strand | N3-A (purple) vs A (green) | N3-A (purple) vs 6mA (orange) | 6mA (orange) vs A (green) |
|---|---|---|---|---|---|
| 46442 | TATCGATGGGC | + | 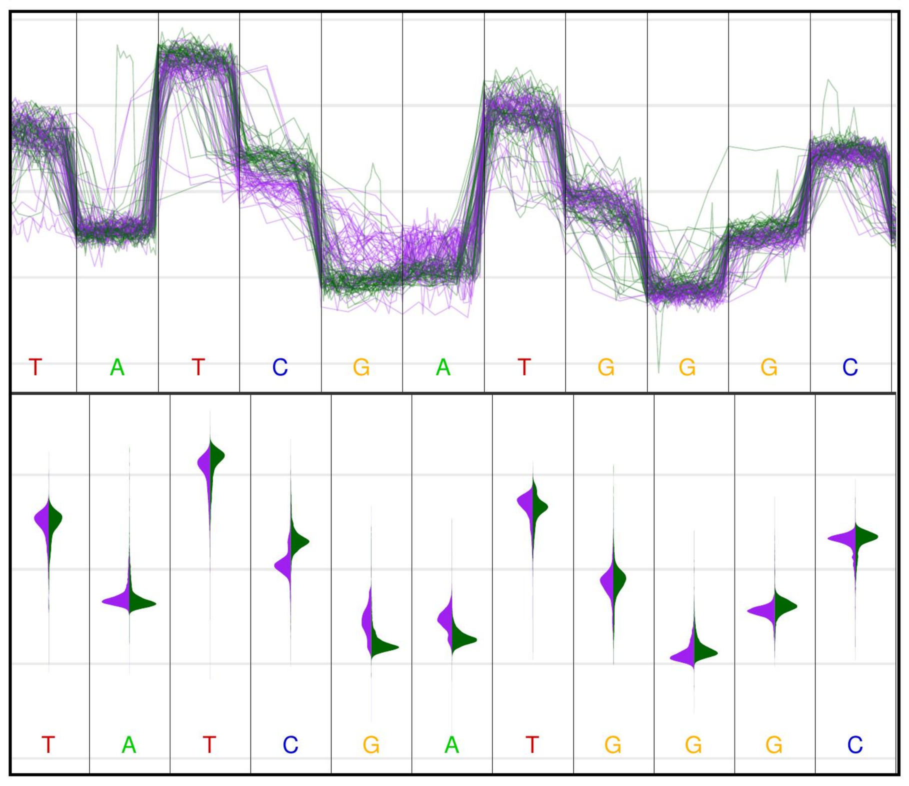 | 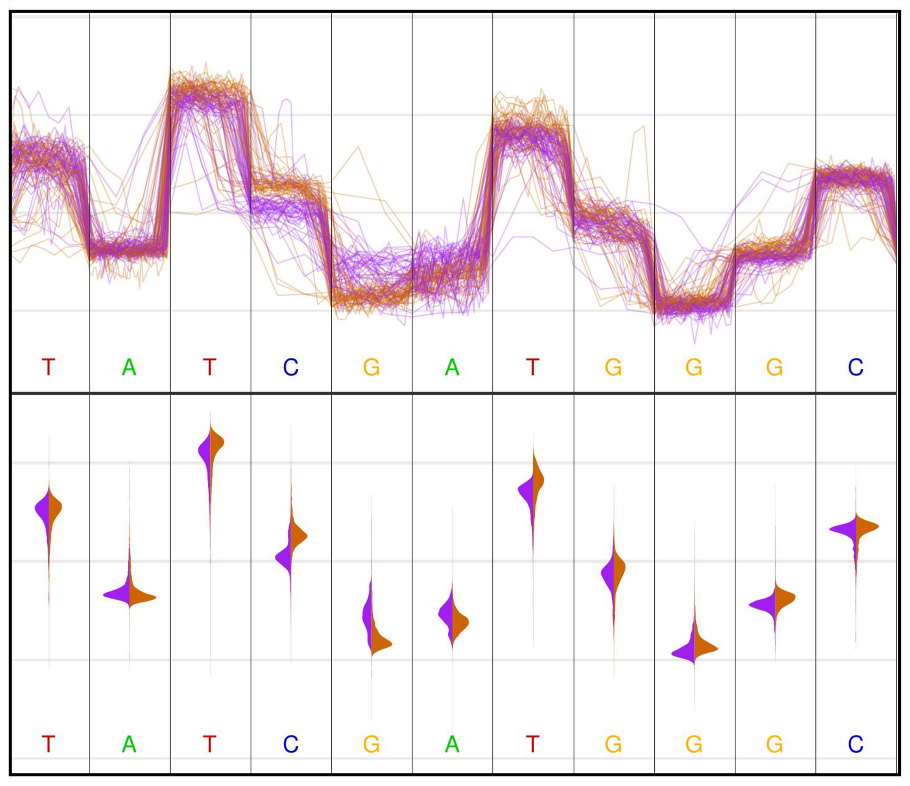 | 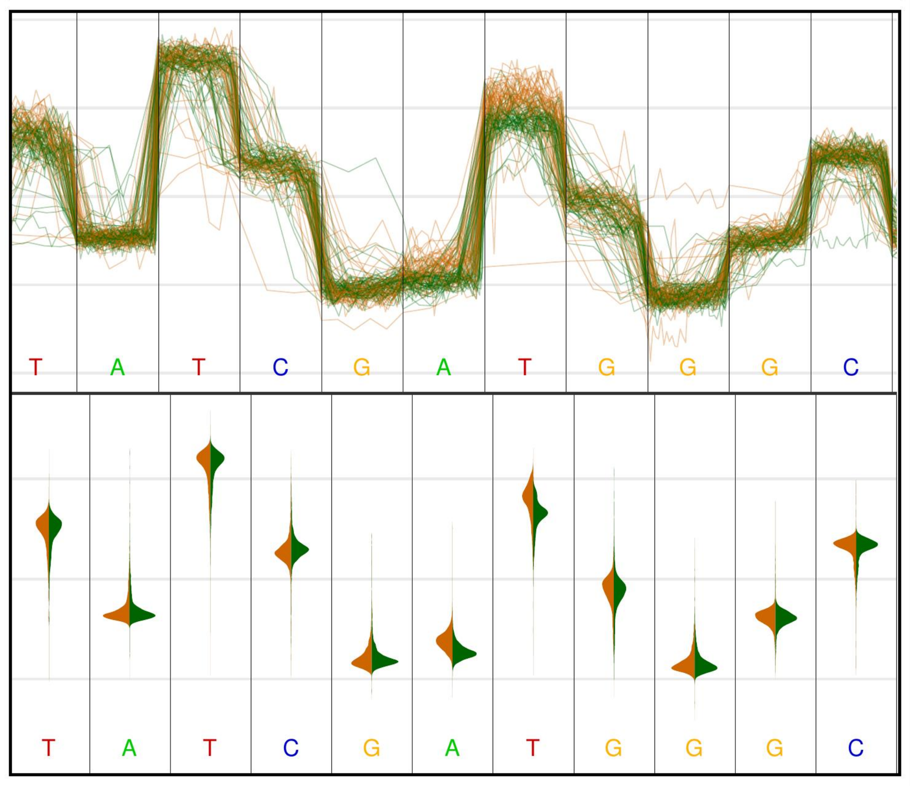 |
| 46439 | CATCGATATGG | - | 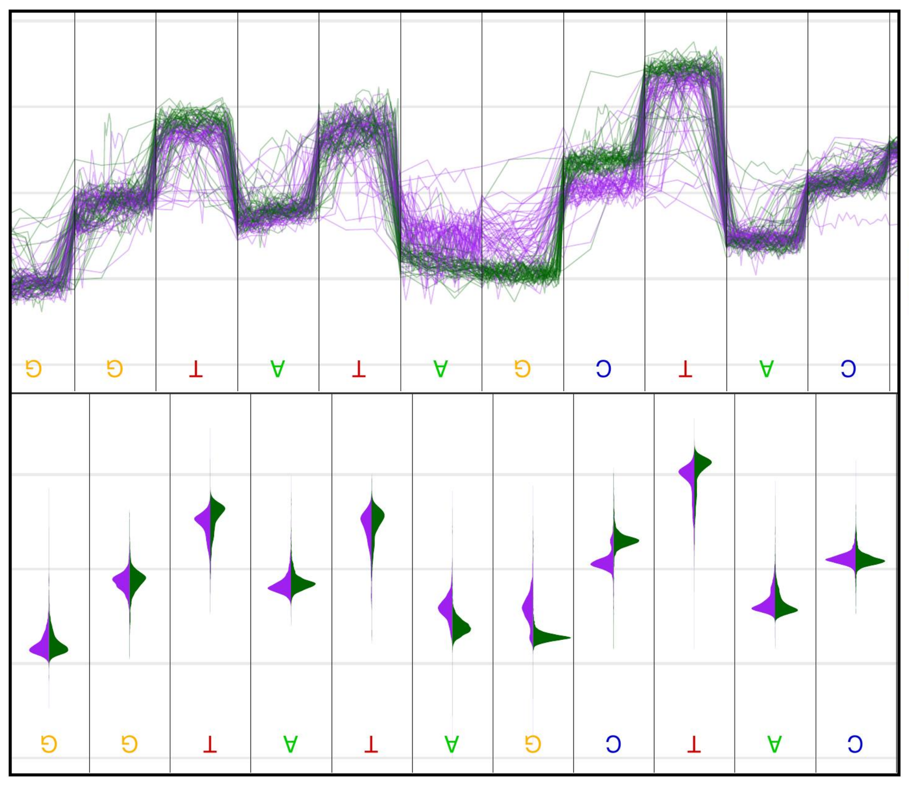 | 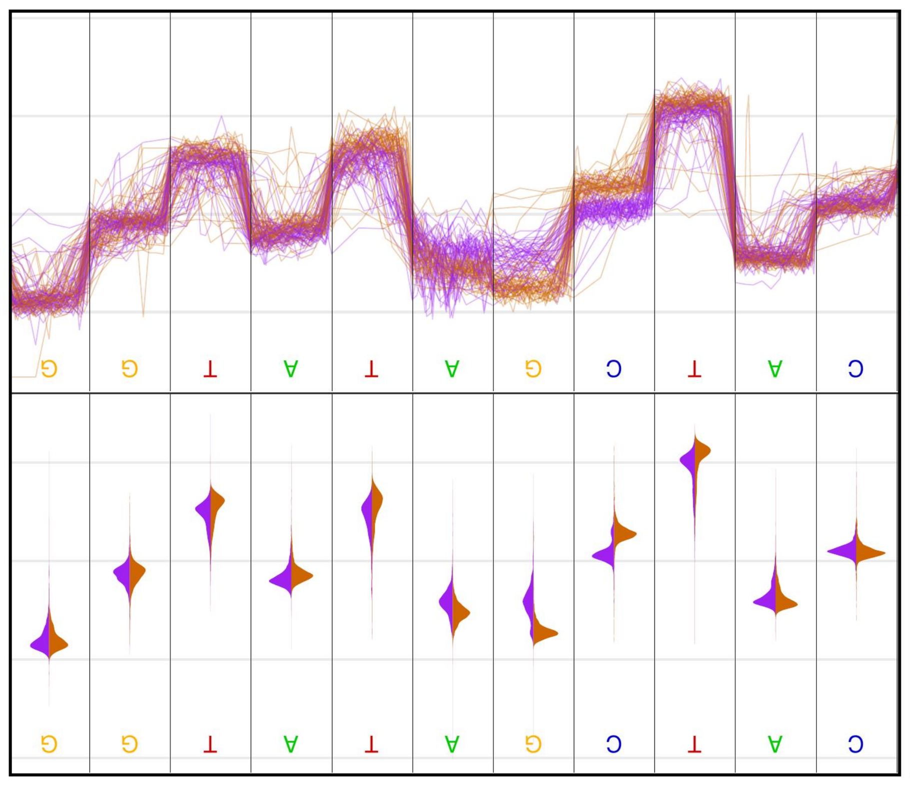 | 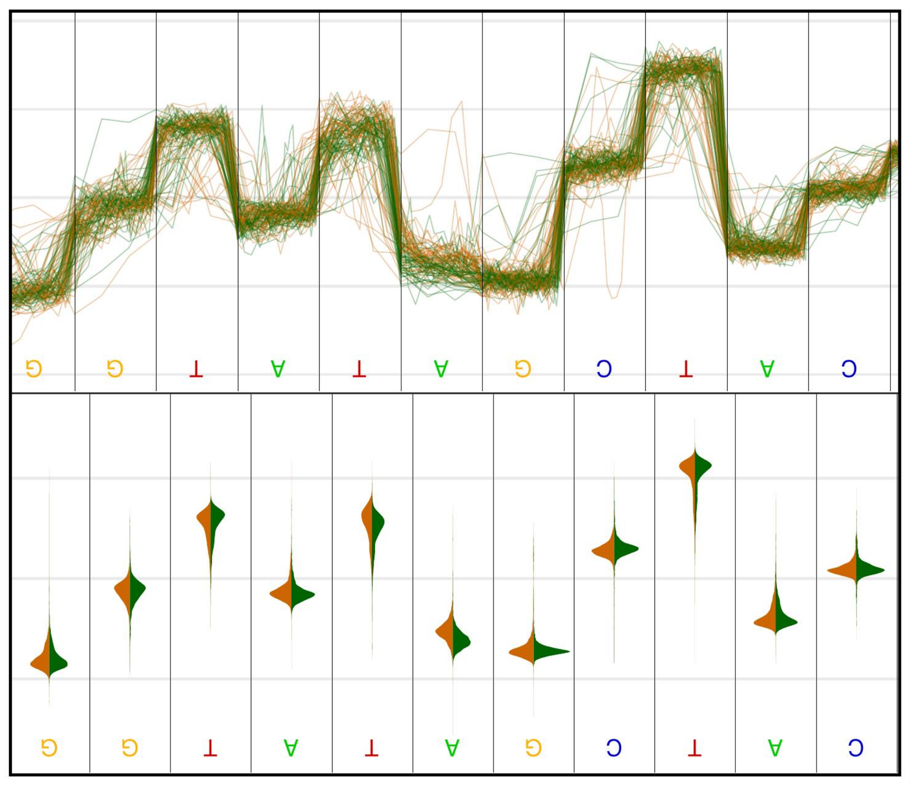 |
| 46598 | CATCGACAGTT | + | 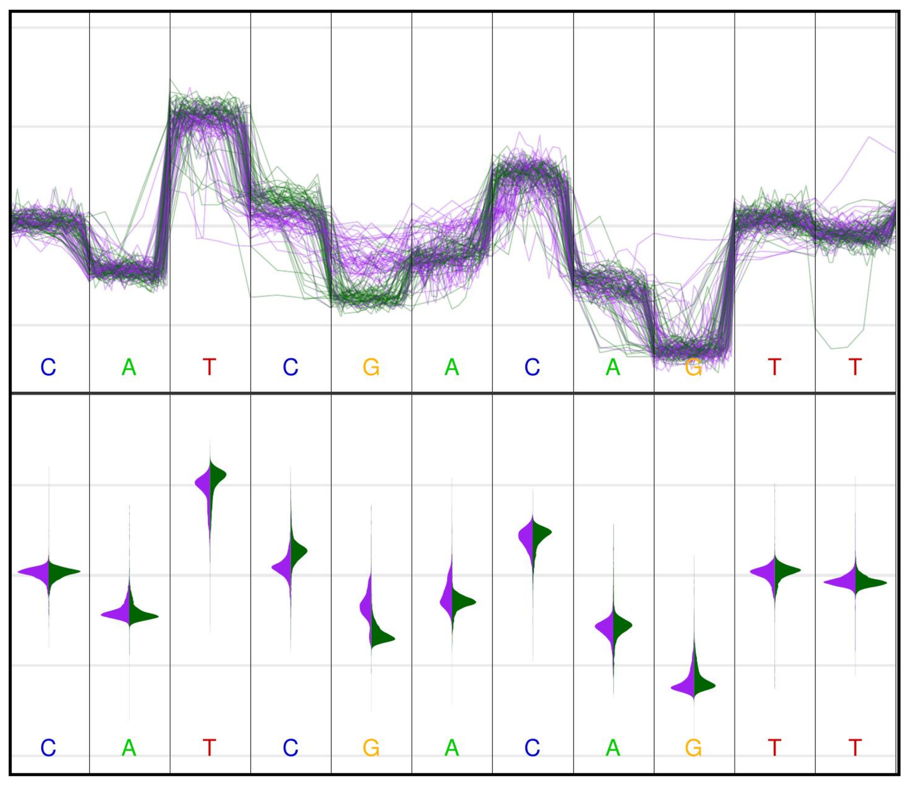 | 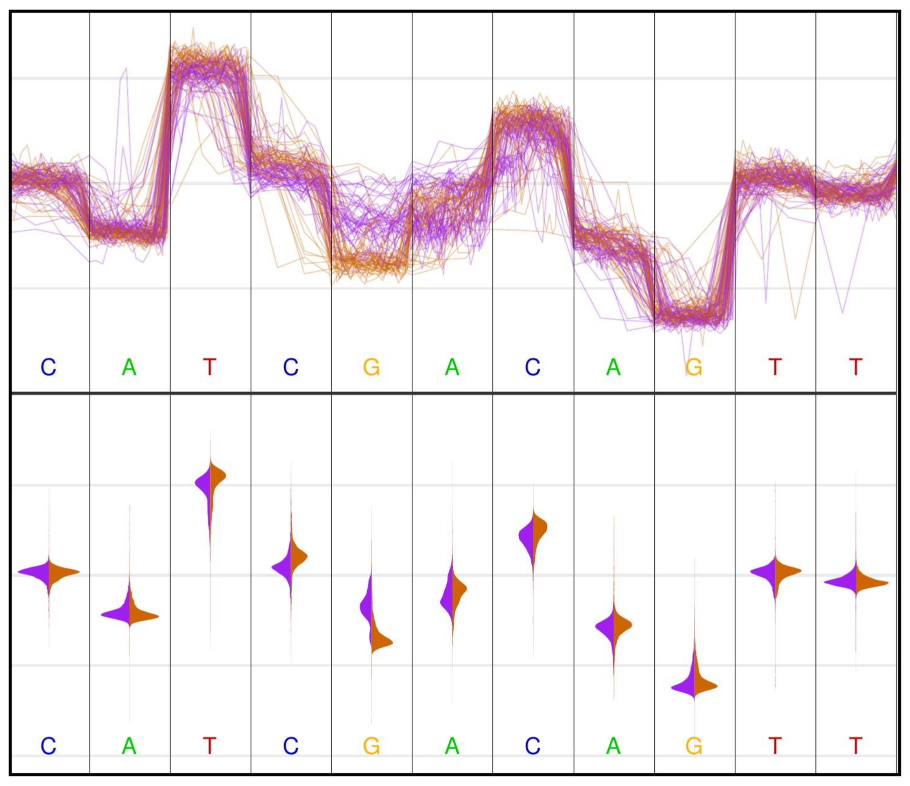 | 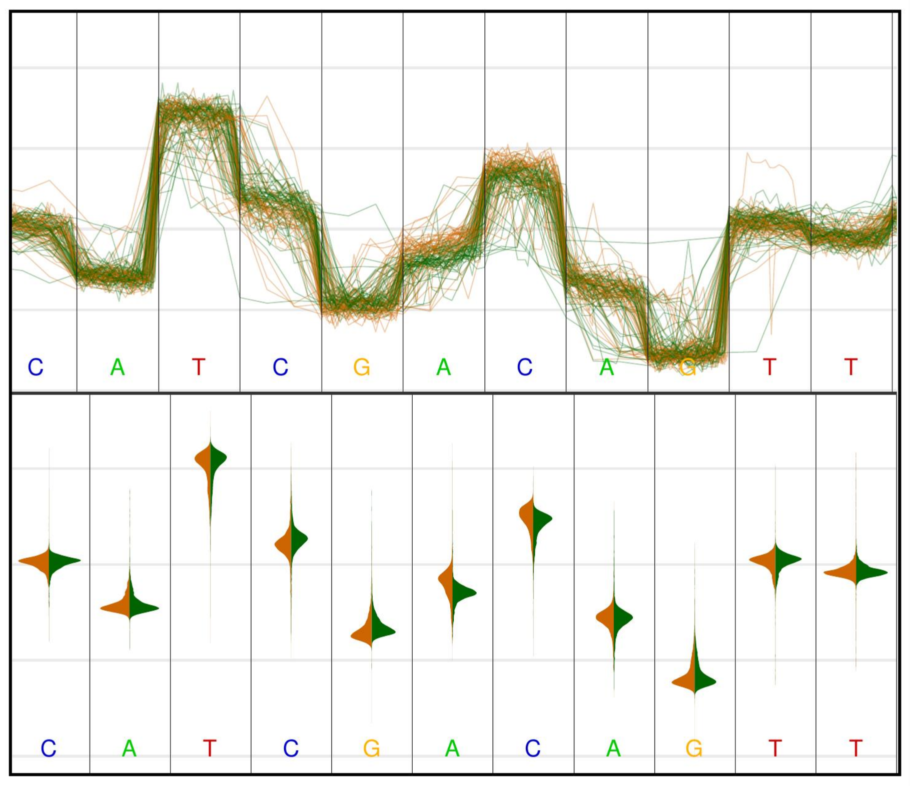 |
| 46595 | TGTCGATGCAG | - | 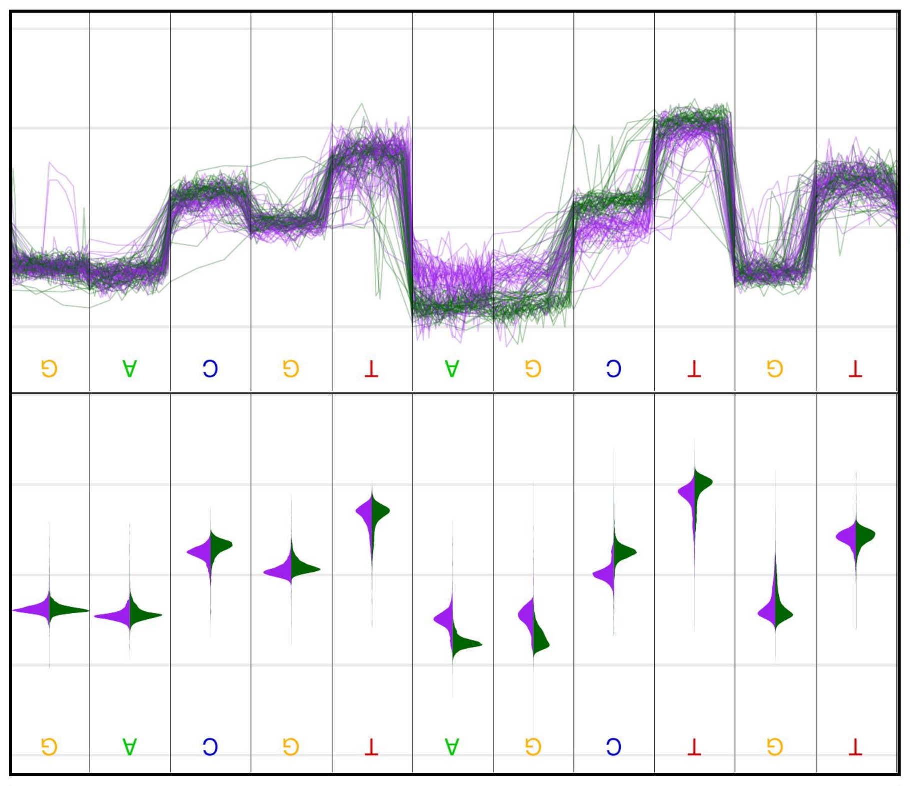 | 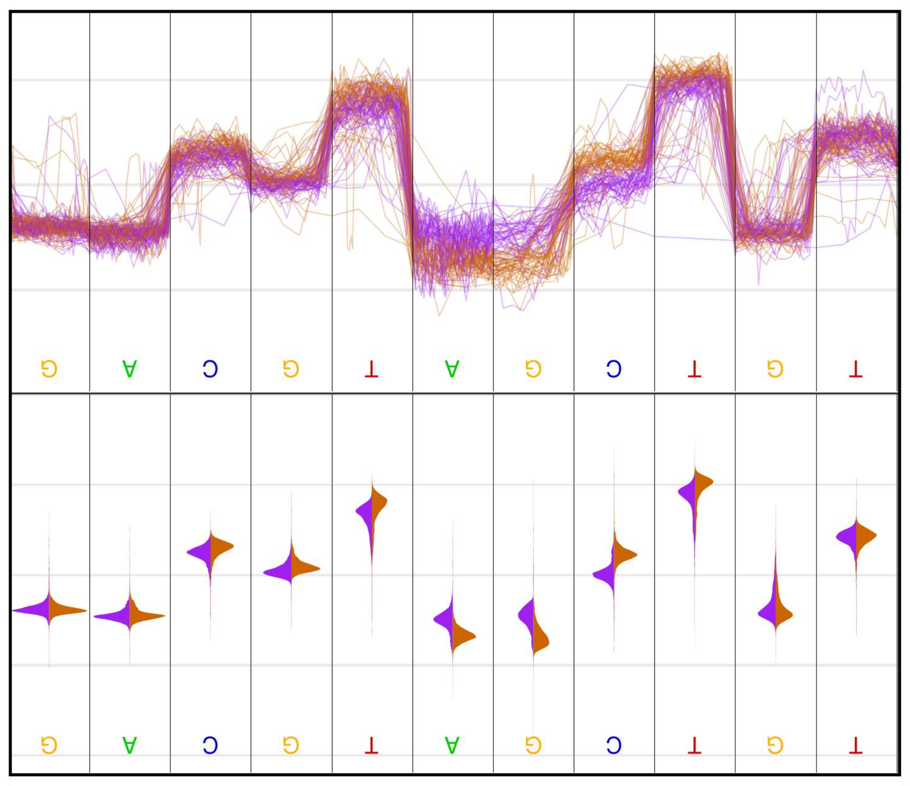 | 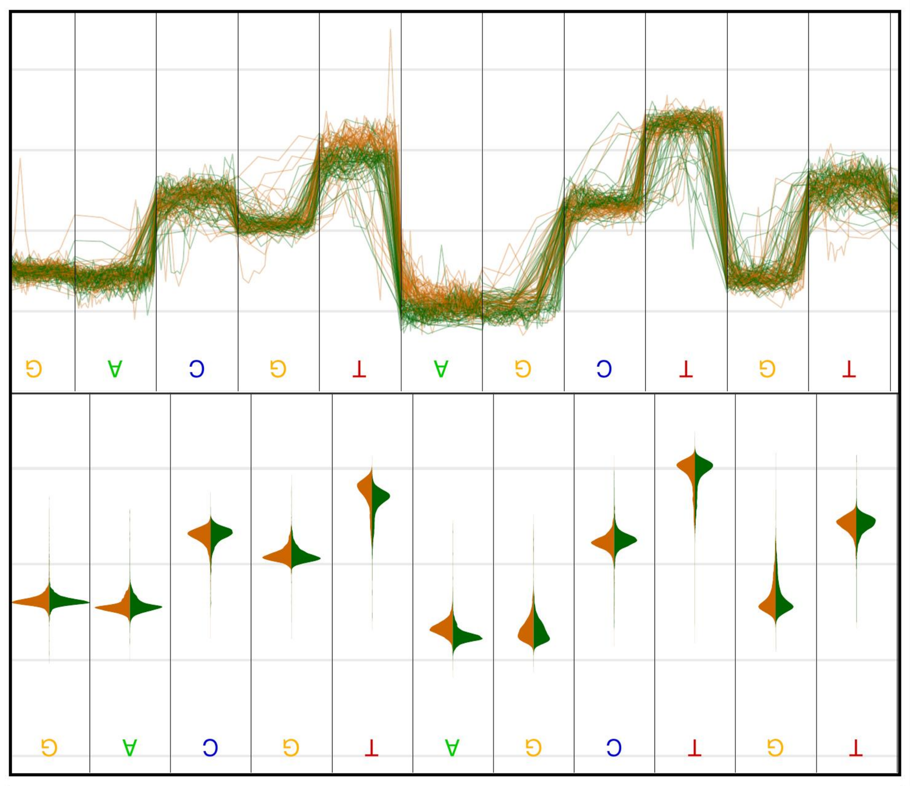 |
| 47421 | GATCGAATATT | + | 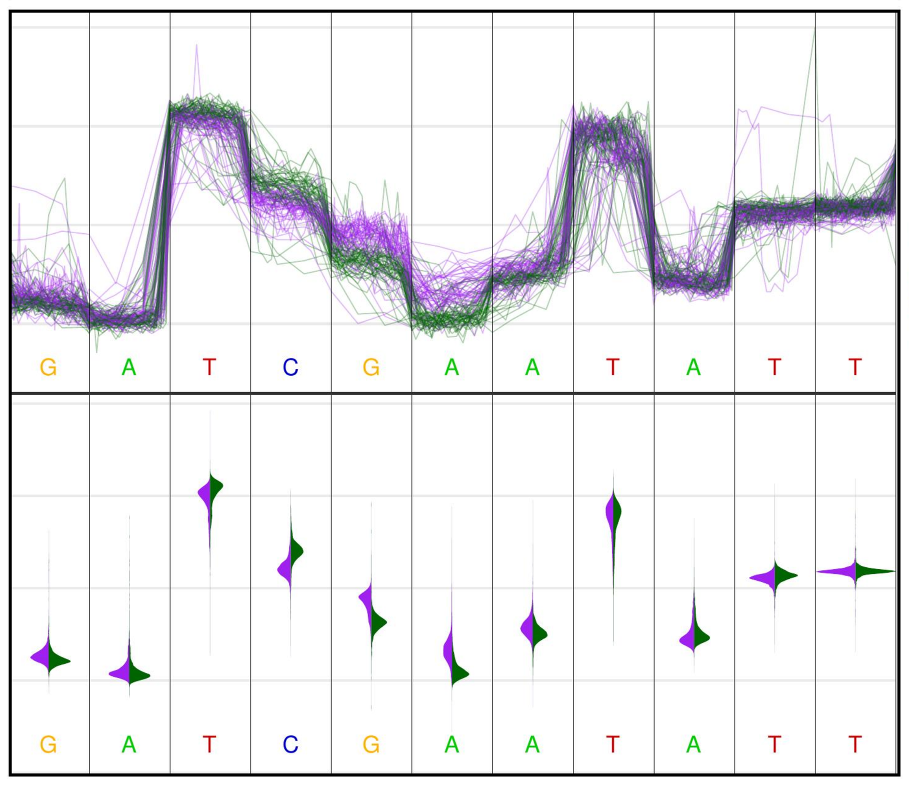 | 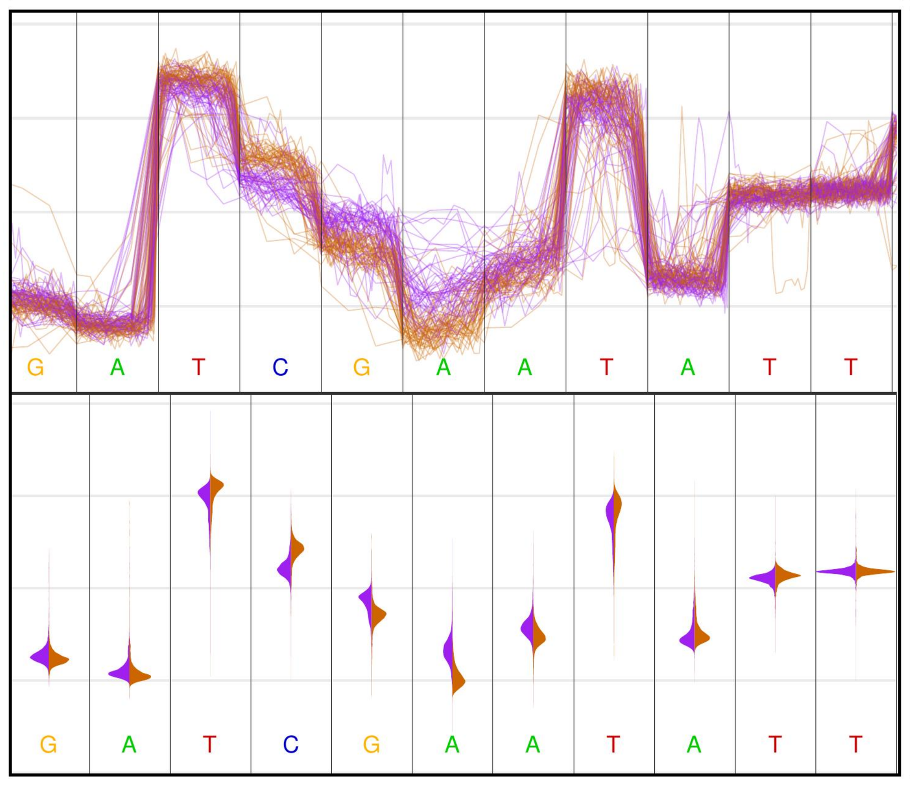 | 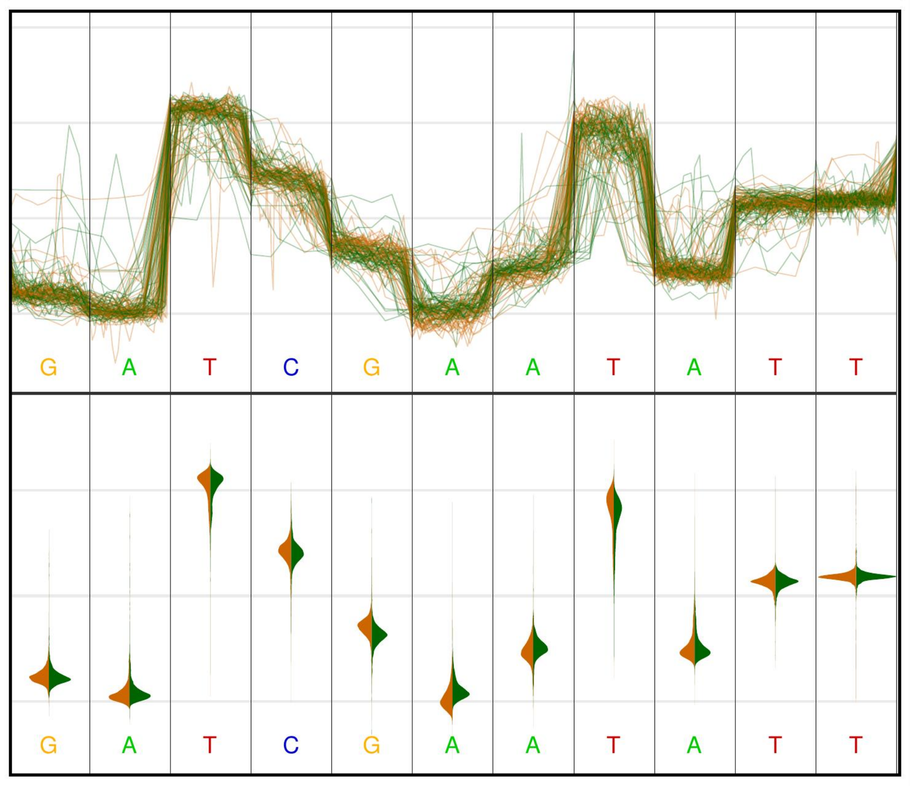 |
| 47418 | ATTCGATCAAG | - | 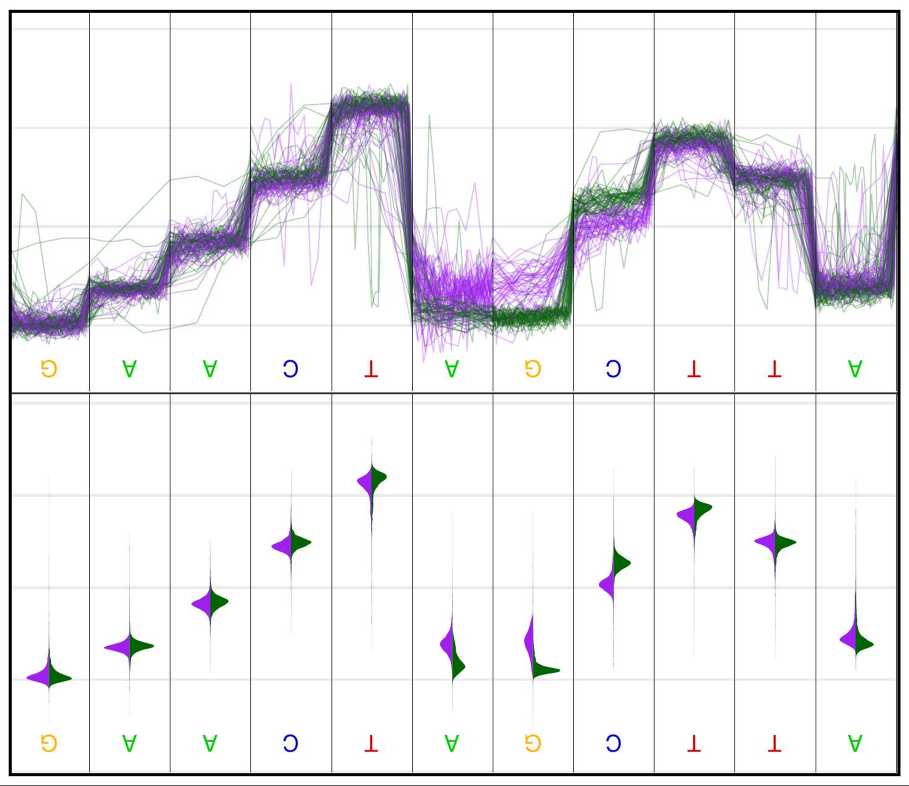 | 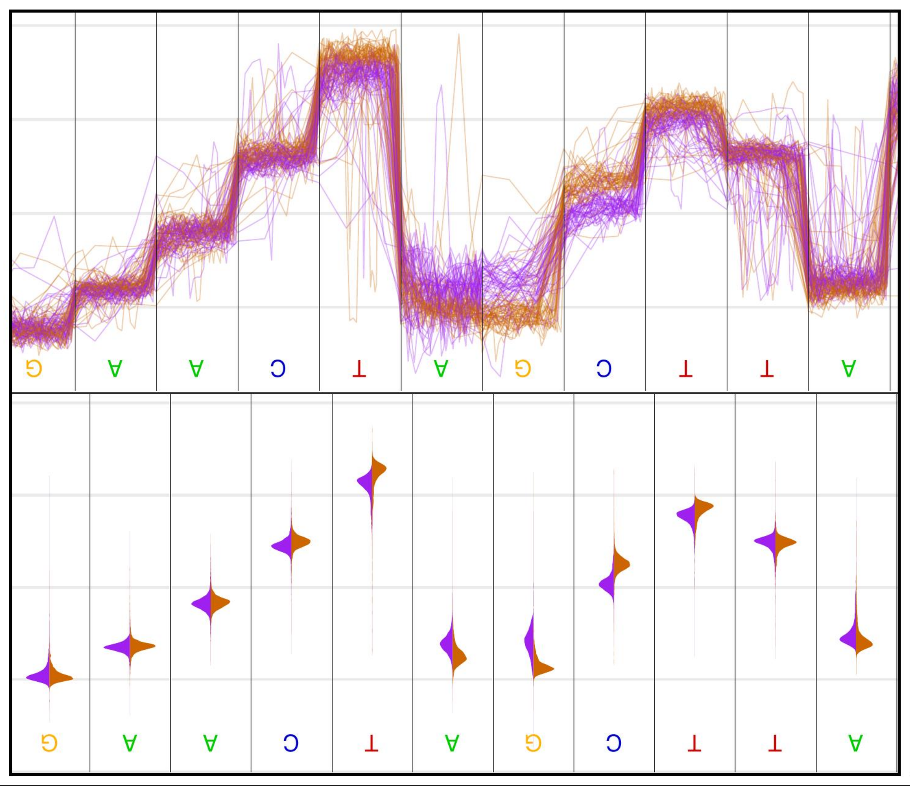 | 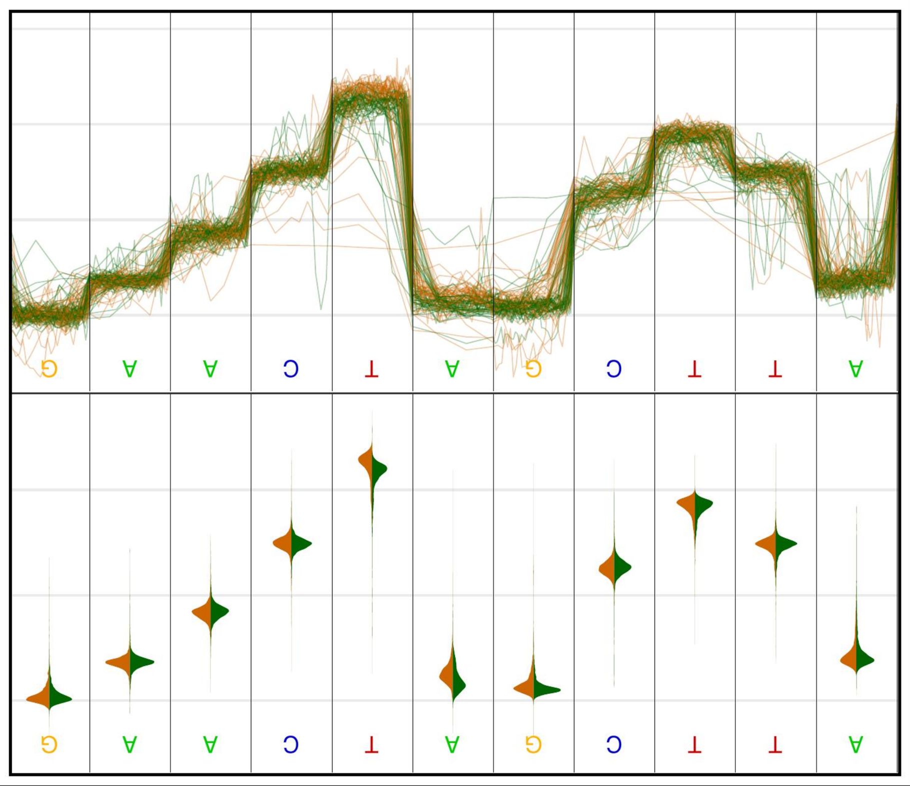 |
